# Multiplexed Deep Visual Proteomics Unveils Spatial Heterogeneity and Rare Endocrine States in Human Adult Pancreatic Islets

**DOI:** 10.1101/2025.04.27.650857

**Authors:** Mariya Mardamshina, Nicolai Dorka, Marvin Thielert, Frida Björklund, Frederic Ballllosera Navarro, Anna Martinez Casals, Ferenc Kovacs, Jocelyn E. Manning Fox, Peter Horvath, Patrick E. MacDonald, Matthias Mann, Emma Lundberg

## Abstract

Pancreatic islets are highly specialized tissue compartments that regulate metabolism, and their dysfunction contributes to diseases such as prediabetes and diabetes. Characterizing islet morphology and cell plasticity is essential for understanding these pathophysiological states, yet high-resolution spatial proteomics remains challenging due to the islets’ small size and cellular complexity. Here, we present multiplexed deep visual proteomics (mxDVP), an approach that integrates high-plex imaging with ultra-sensitive mass spectrometry to achieve deep spatial proteome profiling of defined cell types in tissue sections. Enhanced segmentation, semi-automated annotation, and optimized laser microdissection enable the enrichment of rare endocrine populations that are often overlooked in single-cell analyses. mxDVP achieves deep proteome coverage of >6,000 proteins from as few as 100 cells, including low-abundance transcription factors critical for endocrine cell fate determination. By profiling over 864,000 human pancreatic cells, we identify 12 endocrine subtypes, including polyhormonal hybrids, revealing previously unrecognized islet heterogeneity, metabolic regulation, and cellular adaptability.

## Introduction

The rising global burden of metabolic disorders presents a significant public health challenge, with diabetes cases in the U.S. alone projected to increase by 165%, from 11 million in 2000 to an estimated 29 million by 2050^1^. Additionally, by 2025, over 100 million adults are expected to be affected by prediabetes, with more than 80% unaware of their condition. A key driver of diabetes pathogenesis is insulin resistance coupled with β-cell failure, a process often described as β-cell dedifferentiation and loss of identity, resulting in progenitor-like features^2,3,4^. However, it remains unclear whether these dedifferentiation or potential transdifferentiation events occur in relatively normal islets, highlighting a critical gap in our understanding of pancreatic islet biology.

Pancreatic islets are composed of diverse endocrine populations with distinct morphological and functional attributes essential for metabolic homeostasis. The spatial organization and phenotypic plasticity of these hormone-secreting cells introduce significant challenges in their analysis, particularly in detecting rare endocrine states within the tightly packed islet microenvironment. Traditional single-cell sequencing approaches, while powerful, do not reliably capture polyhormonal cell states, as many studies classify them as doublets or artifacts and exclude them from analysis^5,6^. This limitation obscures the potential biological relevance of these cells and their role in endocrine plasticity. Furthermore, deep proteomic profiling of these cells while maintaining spatial context has been technically challenging, limiting our ability to resolve islet cell heterogeneity at the molecular level.

To address these challenges, we developed multiplexed Deep Visual Proteomics (mxDVP), a comprehensive end-to-end pipeline that integrates high-plex microscopy-based proteomic imaging with ultra-sensitive mass spectrometry-based proteomics. This approach enables spatially resolved, high-depth proteomic profiling within single tissue sections, providing unprecedented molecular insights into pancreatic islet architecture. Key innovations of the mxDVP pipeline include compatibility with multiplexed imaging for accurate visualization of cell types, precise single-cell targeting, semi-automated cell annotation, enhanced segmentation refinement, and automated optimized laser microdissection, facilitating the comprehensive characterization of rare endocrine populations in situ. Notably, Deep Visual Proteomics (DVP) has not previously been applied to single cells in mixed cell populations due to the cutting accuracy demands, nor has it been utilized to analyze adjacent cells in a spatially resolved manner. Moreover, the precise targeting and selection of immediate neighboring cells by laser microdissection has not been performed in such complex and spatially constrained tissues as the islets of Langerhans. The application of mxDVP overcomes these technical limitations, offering a novel method to explore the molecular heterogeneity of pancreatic islets with high spatial precision.

Standard DVP has facilitated significant advancements in the biomedical field, enabling discoveries such as targetable JAK signaling in toxic epidermal necrolysis^7^ and insights into DNA repair and replication stress in signet ring cell carcinoma^8^. Additionally, precise stratification of Hodgkin lymphoma subtypes using standard workflow has revealed novel therapeutic targets, such as IL4-signaling, offering potential strategies for overcoming chemoresistance in the classical subtype^9^. A study by Zheng et al^10^ has demonstrated the power of combining spatial analysis with deep proteomic profiling, revealing targetable metabolic heterogeneity within the immune components and microenvironments of tonsil and colorectal cancers. Building on these insights, mxDVP extends this approach by providing a fully integrated solution for spatially resolved single-cell proteomics, allowing for unprecedented resolution of cell heterogeneity in complex tissue samples. By applying mxDVP to human pancreatic islets, we aim to expand the current understanding of islet cell heterogeneity and its implications for metabolic health and disease. This work establishes a framework for studying rare endocrine states and could inform future research and therapeutic strategies targeting diabetes and related disorders.

## Results

We developed a robust and fully integrated pipeline for mxDVP combining automated multiplexed imaging, laser microdissection (LMD), and ultra sensitive mass spectrometry (Fig.1). This pipeline is optimized for archival formalin-fixed paraffin-embedded (FFPE) tissues, enabling deep single-cell proteomic analysis while preserving spatial context. By leveraging the same tissue section for both imaging and mass spectrometry, we ensure high data fidelity and minimize redundancy.

**Fig. 1:**
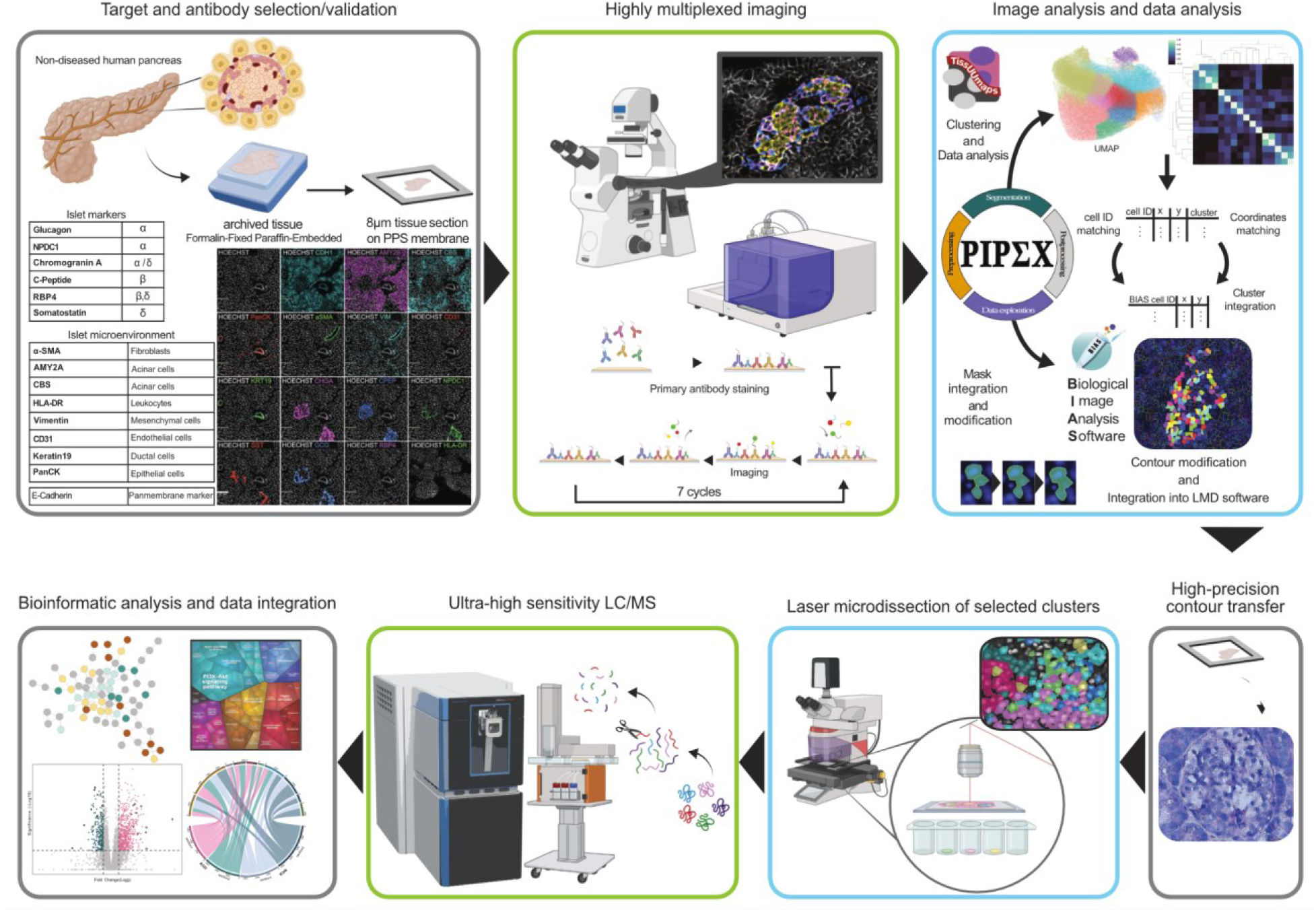
Workflow of integrated mxDVP pipeline. Schematic of the integrated mxDVP pipeline. The pipeline consists of the following key steps (1) development of a customized antibody panel, (2) generation of high-plex spatial proteomic images, (3) image and data analysis through an in-house developed pipeline, (4) introduction of polychromatic stain for high-precision contour transfer, (5) laser-microdissection of selected clusters, (6) MS-based proteomic profiling of isolated cell populations, and (7) integrated data analysis. Grey borders represent novel steps introduced to complete the end-to-end pipeline. Green borders showcase fully automated experimental processes within the pipeline. Cyan borders represent steps in the pipeline that require significant customization.

### Creating an end-to-end automated multiplexed DVP workflow for high-precision proteomics

Pancreatic islets exhibit intricate spatial organization and phenotypic variability, making their profiling particularly challenging. To address this, we designed and validated a 15-plex DNA-barcoded antibody panel tailored for islet cells and their surrounding microenvironment (Extended Data Fig. 1a). This panel enables precise identification of hormone-secreting cells (CHGA, GCG, NPDC1, RBP4, CPEP, SST) alongside exocrine (CBS, AMY2B), immune (HLA-DR), ductal (PanCK, KRT19), and vascular components (VIM, CD31, aSMA), providing a holistic spatial view of pancreatic architecture (Fig. 1).

### Optimizing membrane slide selection for LMD

High-quality laser microdissection relies on optimal membrane slide selection. Through extensive testing, we found that PPS membranes significantly outperformed PEN membranes, offering superior signal-to-noise ratios (PEN: 14.37, PPS: 40.33) and lower autofluorescence. PPS membranes also demonstrated superior durability, and better preservation of tissue integrity throughout the multiplexed staining and cell dissection (Extended Data Fig. 1b).

### Centralized image analysis with PIPΣX

To tackle the complexity of spatial image analysis of the densely packed islet cells, we developed PIPΣX, a custom software that streamlines image segmentation and cell analysis^11^ on the basis of Stardist nuclei segmentation^12^ with membrane refinement. PIPΣX enhances segmentation accuracy, even for irregularly shaped islet cells, by employing the membrane stain CDH1 enabling precise cell boundary definition (Extended Data Fig. 1c). PIPΣX further supports quantification of marker intensity per cell segmentation mask, and clustering, cluster refinement and finally cell population annotation. An initial unsupervised clustering step generates preliminary groups, which are iteratively refined through automated algorithms and manual expert validation to achieve high-confidence cell population annotations (Extended Data Fig. 2a-b).

### Precision refinement for laser microdissection

For enhanced LMD accuracy, PIPΣX was integrated with two other softwares that control the laser microdissection process (Biology Image Analysis Software (BIAS) and Leica LMD software), introducing a post-segmentation adjustment tool to ensure precise isolation of individual cells (Extended Data Fig. 2c).

### Polychromatic staining for precise single-cell isolation

To improve the precision of the contour transfer and single-cell isolation, we implemented a polychromatic staining step using Toluidine Blue after the multiplexed imaging. This approach maintained tissue integrity and did not impact protein expression, ensuring compatibility with downstream proteomic analysis (Extended Data Fig. 2d-e).

### Maximizing proteome depth

Key proteins regulating pancreatic cell function, such as transcription factors, are notoriously challenging to detect due to their low abundance. To overcome this, we optimized cell pooling strategies, progressively increasing the number of cells per sample. We found that the pooling of 250 two-dimensional cell contours (equivalent to the volume of 100 cells) provided the optimal balance between rare cell detection and proteome depth, significantly enhancing the number of unique proteins identified (Extended Data Fig. 2f- h).

### High-resolution spatial profiling unveils diverse endocrine subtypes in normal pancreatic islets

Using our mxDVP pipeline, we generated a detailed spatial profile of endocrine cell populations in normal pancreatic tissue. To ensure reproducibility, two biological replicates with two sections from each donor were included (See Supplementary Table 1). The multiplexed imaging data was processed using the PIPΣX pipeline, enabling precise segmentation and annotation of 864,485 individual cells (Fig. 2a). The dataset included 285,172 acinar cells expressing AMY2B and 139,204 acinar cells expressing CBS and AMY2B. Epithelial (ductal) regions constituted 139,662 cells, while vascular components were represented by 95,925 VIM+ cells, 18,726 αSMA+ cells, and 12,211 CD31+ cells. A subset of 81,620 cells (9.4%) remained unannotated due to limited marker availability. Notably among the annotated cells, 38,753 (5.0%) were identified as islet cells (Fig. 2b).

**Fig. 2:**
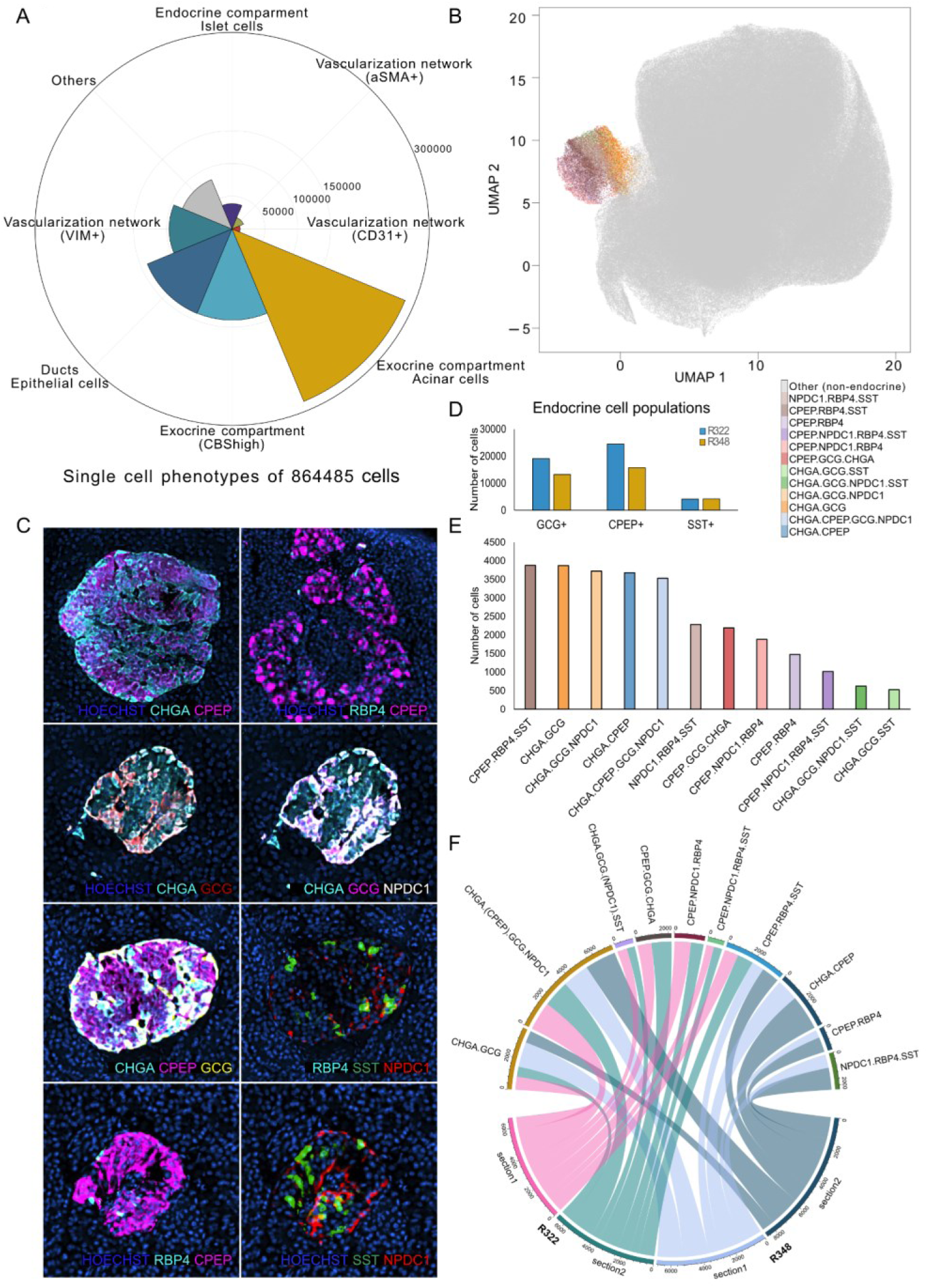
High-plex spatial proteomic profiling reveals distinct endocrine cell populations. a) Rose diagram displaying the number of cells per annotated cell type for the full combined multiplex imaging dataset. b) UMAP representation of the annotated high-plex imaging data of the combined dataset where the endocrine cell population is highlighted and colored by identified subpopulations. c) Representative multiplex images of pancreatic islets with markers distinguishing co-expression patterns identified by K-means clustering. d) Quantification of endocrine cells expressing GCG, CPEP, and SST across donors. The bar plot shows the number of cells per donor. e) Number of cells per identified endocrine cell populations, showcasing both rare and more abundant populations. f) Circos plot comparing the identified endocrine subpopulations between donors and consecutive sections.

Pancreatic endocrine component includes insulin-producing beta cells (β), glucagon-secreting alpha (α) cells, somatostatin-secreting delta cells (δ), pancreatic polypeptide–producing gamma (PP) cells, and ghrelin-secreting epsilon cells (ε). To explore heterogeneity within the endocrine compartment, we applied K-means clustering to the islet cell subset. The clusters were annotated based on their expression profiles, with reference images carefully reviewed for segmentation artifacts and staining imperfections (Extended Data Fig. 3a). In total, we identified 14,491 GCG^+^ cells, 17,689 CPEP^+^ cells, and 8,340 SST^+^ cells (Fig. 2c-d). Established coexpression patterns, such as CHGA and CPEP, CPEP and RBP4, and CHGA and GCG, were readily identified (Fig. 2c; Extended Data Fig. 3b). These patterns were consistently observed in mass spectrometry data, with slight heterogeneity both between and within subtypes, suggesting a gradient-type expression of markers that varies spatially across cell populations (Extended Data Fig. 3c). Additionally, we observed a distinct CPEP, RBP4, and SST co-expressing cell population, within which we detected internal heterogeneity (Extended Data Fig. 3d). Intriguingly, we also detected rare combinations, including SST and NPDC1, as well as triple-marker profiles such as GCG, SST, and NPDC1, underscoring the sensitivity of this approach in identifying rare cell states (Extended Data Fig. 3e). The subclustering revealed 12 endocrine subtypes, characterized by diverse receptor coexpression patterns. The most abundant subtypes included CPEP⁺RBP4⁺SST⁺ (3,875 cells), CHGA⁺GCG⁺ (3,870 cells), and CHGA⁺GCG⁺NPDC1⁺ (3,721 cells), along with CHGA⁺CPEP⁺ (3,673 cells). Rare subtypes, such as CHGA⁺GCG⁺SST⁺ (622 cells) and CHGA⁺GCG⁺NPDC1⁺SST⁺ (527 cells), represented the least abundant populations in the cohort (Fig. 2e).

A comparison across the two donors revealed that the more common subtypes, such as CHGA⁺GCG⁺ and CPEP⁺RBP4⁺SST⁺, were consistently detected across all samples, highlighting their conserved nature. In contrast, rare subtypes, including those with NPDC1 coexpression, were predominantly donor-specific (Fig. 2f). These observations are consistent with patterns identified in publicly available single-cell transcriptomics dataset^13^ (Extended Data Fig. 4a), while also revealing novel hybrid subtypes. While α (GCG⁺GC⁺TTR⁺), β (INS⁺IAPP⁺), and δ (LEPR⁺PRG4⁺SST⁺) cells are traditionally defined by distinct marker expression, accumulating evidence challenges this rigid classification falls short in capturing the underlying biology. Studies have shown that metabolic stressors such as high-fat diet or GLP-1R signaling can induce β-like traits in α-cells, altering their electrophysiology and marker expression^14,15^. Similar plasticity has been observed in type 1 diabetes, where reduced ARX and DNMT1 levels in α-cells coincide with β-like transitions^16^. Our deep proteomic profiling and refined islet cell annotation, identifying both canonical subtypes and hybrid states, including CPEP⁺RBP4⁺SST⁺ and CPEP⁺NPDC1⁺RBP4⁺, highlight the dynamic and adaptive nature of endocrine identity.

### Spatial analysis links islet size to vascularization patterns and polyhormonal cell states

Leveraging the spatial resolution of our multiplexed image dataset, we performed Euclidean distance-based spatial analysis to quantify the proximity of the annotated cell types and their microenvironmental relationships (Fig. 3a). This analysis revealed distinct organizational patterns reflective of functional compartmentalization. Endocrine cell populations exhibited closer spatial distances among themselves compared to their proximity to exocrine or ductal cells, underscoring their cohesive niche organization into islets of Langerhans. Similarly, exocrine cell types displayed lower Euclidean distances within their compartment, highlighting their spatial clustering and potential cooperative roles in digestive processes. In contrast, vascular and ductal compartments were spatially segregated from other cell types, indicating distinct structural organization within the tissue (Extended Data Fig. 4b).

**Fig. 3:**
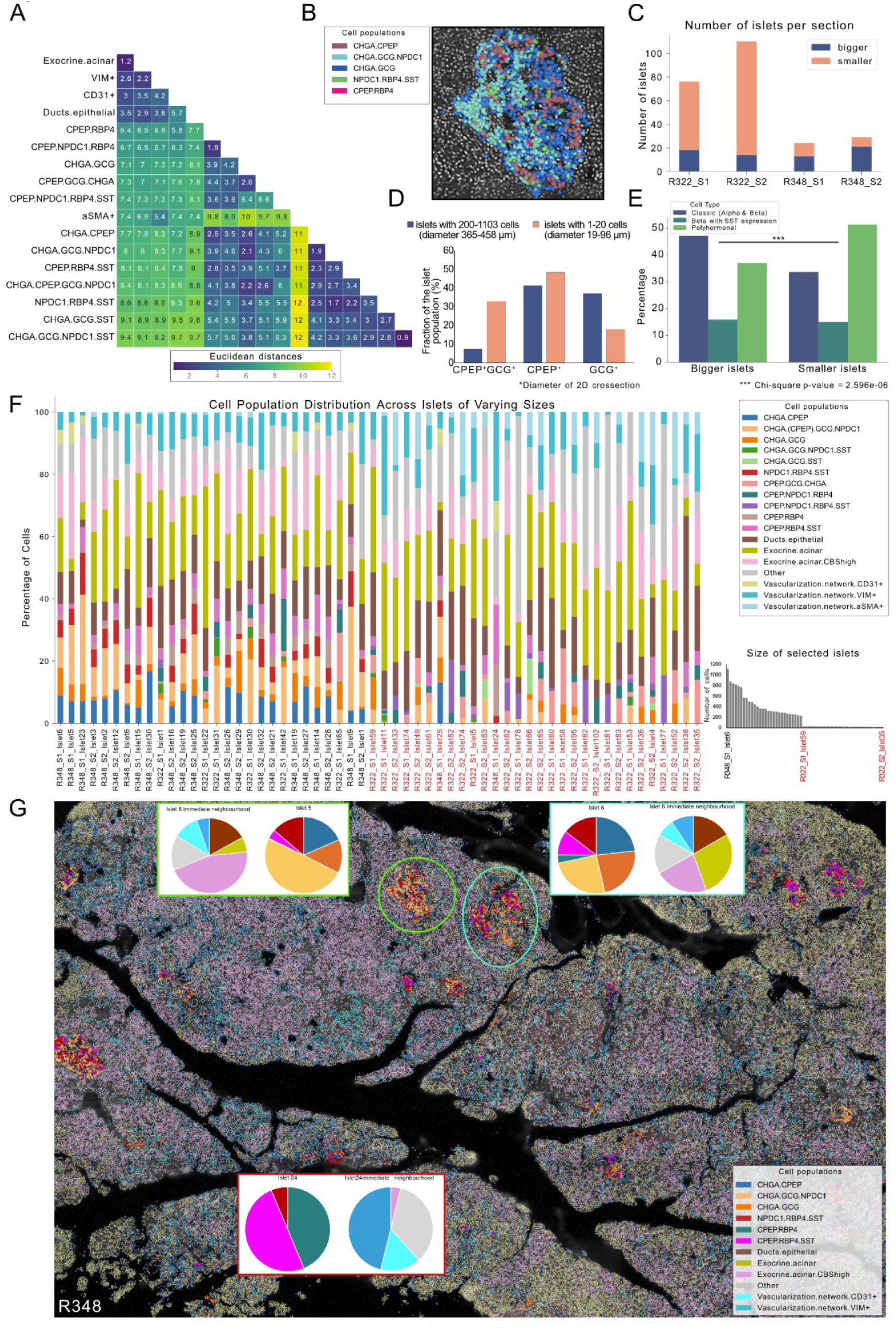
Islet-centered spatial analysis unveils that islet size correlates with cell type composition of islets and the vascular bed. a) Heatmap displaying the Euclidean distances between all annotated cell populations, quantifying the spatial relationships in the tissue microenvironment. b) Example image of observed islet cell type compartmentalization.

Cell segmentation masks are colored by the annotated cell type and overlayed on the nuclear stain image. c) Quantification of the number of islets per sample, showcasing the difference in islet size between sections. d) Barplot demonstrating proportions of GCG-, CPEP-, and GCG/CPEP-expressing cell populations in small islets (fewer than 20 cells) and large islets (more than 200 cells). e) Comparison of cellular composition in different-sized islets. Statistical significance was determined by the Chi-square test (p-value = 2.595e-6). f) Proportions of annotated islet cell populations and immediate surrounding vasculature for selected different-sized islets across all samples. g) Overview of one tissue section (R348_S1) with annotated cell populations overlaid on the nuclear stain image. Islets of different sizes are highlighted, and the cellular composition of the islets and their immediate neighborhoods are visualized by pie charts.

Within the vascular compartment, we observed a correlation between marker expression and vessel caliber. Specifically, αSMA⁺ cells were predominantly associated with larger-caliber vessels, whereas CD31⁺ endothelial cells were primarily localized to smaller vessels (Extended Data Fig. 4c). Spatial analysis further supported this distinction, as αSMA⁺ cells were positioned at greater distances from endocrine cell clusters, suggesting potential spatial segregation between these populations. Compartmentalization was also evident within pancreatic islets. Similar to the patterns reported by Bosco et al.^17^, α-cells, identified by GCG expression, were predominantly localized to the islet periphery, whereas β-cells, marked by CPEP expression, were concentrated toward the islet center (Fig. 3b; Extended Data Fig. 4d). Despite this spatial partitioning, α- and β-cells remained in close proximity, as indicated by low Euclidean distances, likely facilitating coordinated interactions crucial for metabolic regulation^18,19^. Rare cell populations exhibited unique spatial distribution patterns. Some formed localized clusters, whereas others, including cells expressing NPDC1, RBP4, and SST, were sparsely dispersed across the islets (Extended Data Fig. 4e). These sparsely distributed populations exhibited higher Euclidean distances relative to other islet cell types, suggesting distinct microenvironmental roles.

Building on these spatial observations, we next examined the relationship between islet size (i.e. two-dimensional cross-section) and cell type composition, both across donors and within consecutive tissue sections from the same donor (Fig. 3c). First, we examined the relationship between islet size and cellular composition by assessing the distribution of GCG-, CPEP-, and GCG/CPEP-expressing cells. Consistent with the findings of Lehrstrand et al.^20^, we found that GCG-expressing cells were predominantly localized within larger islets, whereas smaller islets contained a higher proportion of CPEP-expressing cells. Notably, the highest percentage of GCG/CPEP co-expressing cells was observed in smaller islets. To further quantify this relationship, we analyzed islet size in relation to endocrine cell populations, revealing a significant association between the two variables (Chi-square test, Fig. 3e). This structural heterogeneity within the pancreatic microenvironment prompted us to investigate the peri-islet niche by expanding analysis regions to 1.5 times the diameter of each islet across our dataset (Extended Data Fig. 4f). For this comparison, islets were categorized as small (<20 cells) or large (>200 cells) (Fig. 3f). It is important to note that the size classification was based on two-dimensional tissue sections, and some small islets may represent cross-sections of the top of larger islets. Larger islets primarily consisted of classical α- and β-cells, with CHGA⁺ and CPEP⁺ cells predominantly localized within them (Fig. 3g). In contrast, smaller islets were enriched in α- cells and polyhormonal populations, often displaying a relatively uniform representation of specific endocrine subtypes. This suggests that smaller islets may reflect localized environmental demands or dynamic shifts in endocrine cell mass. We next examined the peri-islet microenvironment and its relationship to islet size. Large islets exhibited lower vascular representation, particularly of αSMA⁺ cells, and a higher proportion of acinar cells with elevated CBS expression. This pattern may indicate hypoxic states due to sparse vascularization, as previously suggested^22^. In contrast, small islets displayed increased vascular representation, including larger vessels containing αSMA⁺ and VIM⁺ cells. This enhanced vascularization could be critical for meeting the metabolic and oxygen demands of growing islets, providing a structural framework for nutrient and oxygen delivery^23^.

### Deep proteomic characterization of endocrine cell populations

To uncover the molecular diversity of pancreatic endocrine cells, including rare polyhormonal states, we leveraged the mxDVP pipeline for deep proteomic profiling. Using laser microdissection, we precisely isolated subpopulations from the same multiplexed tissue sections, integrating AI-driven annotation and automated cell targeting (Fig. 4a; Extended Data Fig. 5a). Each subpopulation was processed independently across multiple donors and tissue sections to ensure robust representation.

**Fig. 4:**
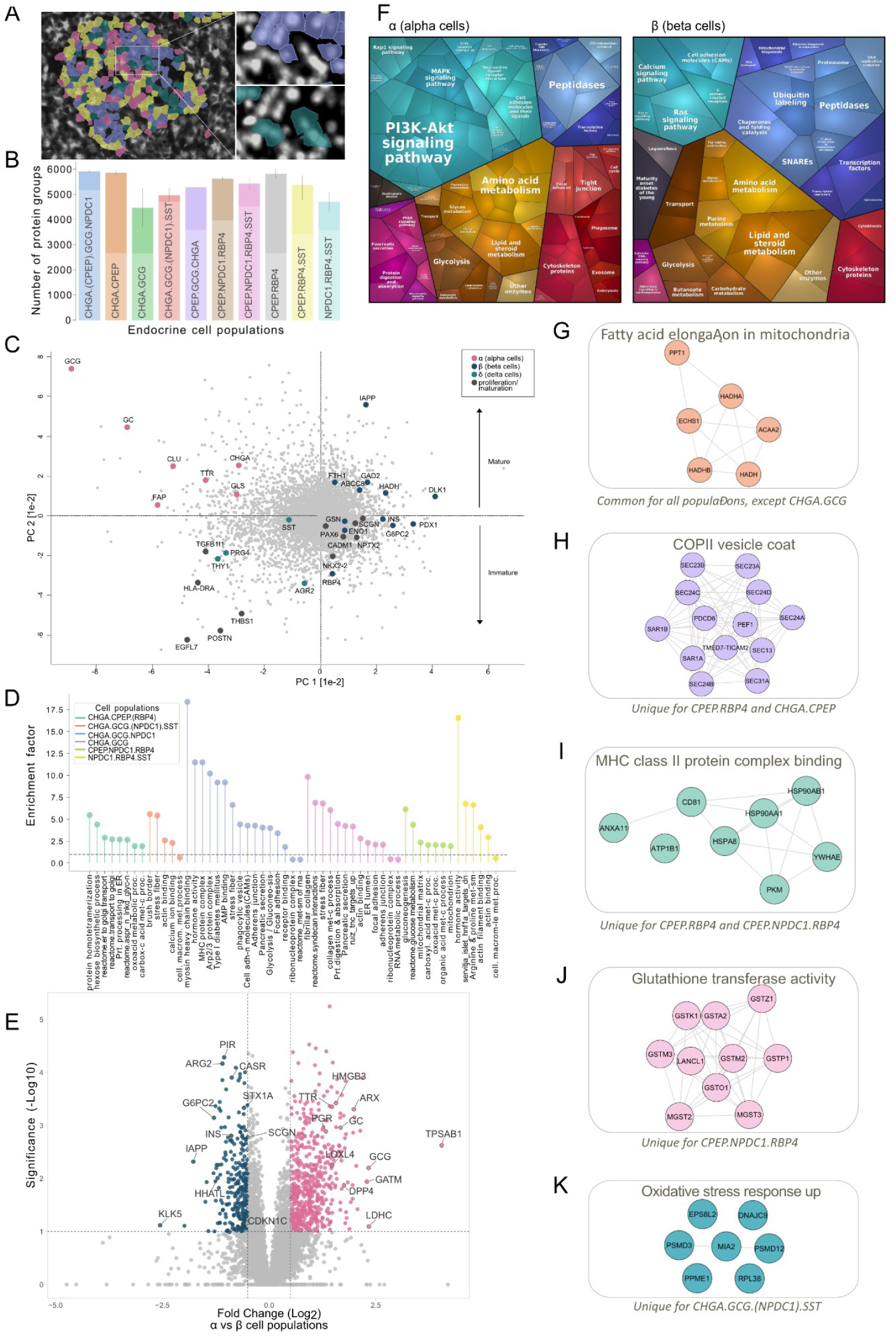
Integrated mxDVP pipeline allows for proteomic characterization of distinctive islet cell populations. a) Example image of cell segmentation masks colored by integrated AI-driven annotation of islet cell type showcasing the precise cell populations isolated by laser-microdissection and analyzed by MS. b) Number of protein groups identified by MS for each isolated endocrine cell population. c) Principal component analysis of the entire cohort after outlier removal with protein markers in α-cells (pink), β-cells (blue), δ-cells (turquoise), and proliferating/maturing cells (dark gray) highlighted. d) One-dimensional enrichment analysis of cellular processes associated with enriched proteins for each endocrine cell population (FDR < 0.05). e) Volcano plot comparing α-cells (pink) and β-cells (blue) proteomic profiles. Welch t-test (FDR < 0.05) revealed significant subtype-specific protein signatures. f) Proteomaps^21^ pathway enrichment analysis illustrating functional processes enriched in α-cells (left) and β-cells (right). Each polygon represents a KEGG pathway, with size correlating with the ratio between the groups. This visualization highlights distinct functional processes between the two endocrine cell types. g-k) Protein-protein interaction networks of the endocrine subpopulations were generated based on the String database, and colored based on enriched pathways (Fisher exact test Benjamini-Hochberg FDR q-value < 0.02).

Across both donors, we identified about 6,000 unique proteins, revealing distinct and deep proteomic signatures among classical islet cell types (Fig. 4b; Extended Data Fig. 5b-c). Principal component analysis (PCA) showed clear segregation between α- and β-cells along PC1, while PC2 correlated with cellular maturity (Fig. 4c). For instance, α-cells were enriched for GCG, GC, CLU, TTR, CHGA, GLS, and FAP, reinforcing their roles in glucagon secretion and metabolic regulation. Delta (δ) cells, marked by SST, PRG4, AGR2, and THY1, suggested a proliferative, maturing population^24^. β-cells exhibited heterogeneity, with mature cells expressing IAPP, HADH, DLK1, FTH1, and ABCC8, while immature β-cells were distinguished by PDX1, ENO1, G6PC2, and RBP4^25,26^. Notably, markers linked to β-cell differentiation, PAX6, NKX2-2, and CADM1, were enriched in immature populations, suggesting ongoing maturation or potential cycling between insulin-high and insulin-low states ^27–29^.

To explore functional differences across cell types, we performed one-dimensional enrichment analysis (Fig. 4d). β-cells were enriched in metabolic processes, including hexose biosynthesis, carboxylic acid metabolism, and ER-to-Golgi transport. α-cells, in contrast, were associated with hormone activity, focal adhesion, and ER lumen function^30,31^. A comparative analysis (Welch t-test; FDR <0.05) identified hundreds of subtype-specific proteins, with α-cell markers (GCG, TTR, GC, ARX, LOXL4) and β-cell markers (INS, IAPP, PIR, G6PC2, SCGN) showing distinct clustering (Fig. 4e). Pathway analysis further revealed PI3K-Akt signaling, amino acid metabolism, and PPAR signaling in α-cells^32^, while calcium signaling, Ras signaling, and lipid metabolism were enriched in β-cells ^33,34^ (Fig. 4f, Extended Data Fig. 5d).

Distinct metabolic demands were observed among endocrine subtypes: β- and δ-cells relied on fatty acid oxidation for sustained ATP production, whereas α-cells prioritized glucose sensing and glucagon secretion (Fig. 4g). Within β-cells, CPEP-expressing cells, identified as insulin-high cells based on our mass spectrometry data, exhibited enhanced COPII vesicle transport, a pathway critical for efficient proinsulin processing and secretion (Fig. 4h). This suggests that CPEP- expressing cells may upregulate COPII transport under metabolic stress to sustain insulin output^35^. Beyond metabolic adaptation, pancreatic islet cells co-expressing CPEP and RBP4 exhibited enrichment in the MHC class II protein complex binding pathway, suggesting a potential link between metabolic regulation and immune modulation (Fig. 4i). While C-peptide itself is not traditionally known for direct involvement in immune presentation, its presence alongside RBP4 may indicate metabolic stress or inflammation-associated changes in antigen processing. RBP4 is essential for retinol transport, and elevated levels have been linked to insulin resistance, raising the possibility that RBP4-expressing islet cells contribute to local immune signaling^36^. Although their direct role in diabetes pathogenesis remains unclear, the enrichment of MHC class II components suggests a potential function in antigen presentation within the islet microenvironment, warranting further investigation.

Beyond classical islet cell types, we identified rare populations exhibiting unique metabolic and stress-adaptive signatures, including gluconeogenesis, oxidative stress responses, and glutathione transferase activity (Fig. 4j, k)^37^. Notably, elevated glutathione synthesis has been linked to stressed β-cell states^38^, and shifts in arginine/proline metabolism have been associated with cellular plasticity and stress adaptation, mechanisms that may help preserve islet function and prevent apoptosis^39^.

Together, these findings demonstrate how deep proteomic profiling unveils endocrine cell heterogeneity, capturing metabolic demands, signaling networks, and stress adaptation mechanisms that shape pancreatic islet function.

### Exploring the spatial proteome of rare and bihormonal endocrine cell populations

Next, we focused on rare endocrine cell populations and started by investigating proteomic differences between classical subtypes and bi-hormonal cells. An ANOVA test (p = 0.05) followed by a PostHoc Tukey’s HSD test (FDR = 0.05) identified hundreds of upregulated proteins distinguishing these populations (Fig. 5a; Supplementary Table 2). Markers characteristic of the β-cells, including IAPP, G6PC2, SCGN, PICK1, and HADH, were detected in both classical β- and bi- hormonal (beta with SST expression) cells. Additionally, bi-hormonal cells exhibited co-expression of markers commonly shared between β- and δ-cells, such as CADM1, PDX1, and PCSK1^40^. PDX1, a master regulator of pancreatic development and β-cell function, plays a crucial role in both the formation and maintenance of insulin-producing β-cells, including their transdifferentiation from non-β-cells^41,42^.

**Fig. 5:**
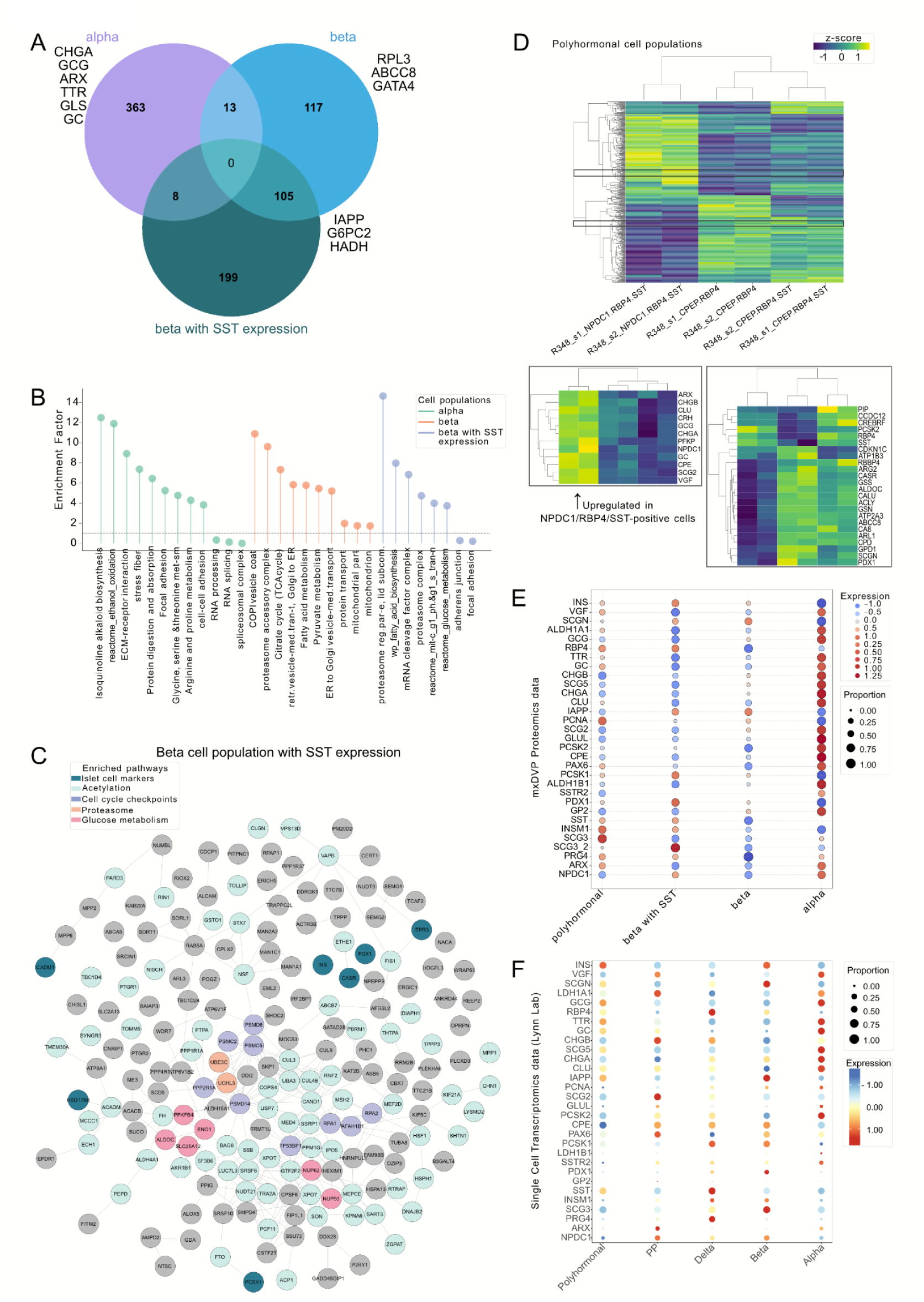
Deep proteomic profiling of bihormonal and polyhormonal cells uncovers rare dynamic endocrine cell states. a) Venn diagram displaying proteomic differences between α-cells, β-cells, and bihormonal cells. Statistical significance of expression differences was determined by ANOVA test (p-value > 0.05) followed by PostHoc Tukey’s HSD test (FDR > 0.05). b) Enrichment analysis of cellular processes associated with enriched proteins, comparing cellular processes enriched in α-cells, β-cells, and bihormonal beta cells with SST expression. c) Protein-protein interaction networks of the bihormonal beta cell population with SST expression were generated based on the String database, and colored based on enriched pathways (Fisher exact test Benjamini-Hochberg FDR q-value < 0.02). d) Heatmap showing all differentially expressed proteins identified by mass spectrometry in selected RBP4-positive cell populations across the cohort (top). The first highlighted region (bottom left) illustrates representative α-cell-associated markers. The second highlighted region displays representative β-cell- and δ-cell-associated markers, along with several differentially expressed prohormones. e) Dot-plot of mxDVP MS-based proteomic data visualizing the protein expression and proportion across polyhormonal, bihormonal, β-cells, and α-cells for key islet cell markers. f) Dot-plot of single-cell transcriptomic data from a publicly available dataset^50^ showing the gene expression and proportion across polyhormonal, bihormonal, β-cells, and α-cells for key islet cell markers.

Enrichment analysis revealed a strong metabolic signature in both classical β- and bi-hormonal subtypes, with pathways involved in the TCA cycle, fatty acid metabolism, and glucose metabolism (Fig. 5b). Notably, bi-hormonal cells also displayed enrichment for pathways associated with cell cycle regulation, mitotic G1 and G1-to-S transition, proteasome complexes, acetylation, and adhesion (Fig. 5c). Cell adhesion, which plays a key role in insulin processing and exocytosis, has also been found to be enriched in bi-hormonal cell populations in zebrafish^43^. Furthermore, acetylation plays a critical role in pancreatic progenitor differentiation into mature β-cells by modulating key transcription factors such as PDX1 and MAFA, which govern insulin production and β-cell identity^44^. Notably, β-cells undergoing cell cycle entry often exhibit transient downregulation of key maturation markers, resulting in a less differentiated state that may temporarily reduce insulin production^45^. Given that bi-hormonal cells display both cell cycle activity and shared β- and δ-cell characteristics, their identity may reflect an intermediate or transdifferentiating state within the dynamic landscape of β-cell plasticity.

Expanding our analysis from bi-hormonal to the rarer polyhormonal cell populations, we identified subtype-specific markers whose expression patterns aligned with multiplexed imaging data (Extended Data Fig. 5e). Among 23 overlapping proteins detected in NPDC1-positive polyhormonal cells, we observed a hybrid signature, with co-expression of α-cell markers (GCG, CHGA, PGR, MAOB) and β-cell markers (MAOA, ABCB10, VIL1) (Extended Data Fig. 5f). Interestingly, these polyhormonal cells also expressed DPP4, typically described as an α-cell marker in humans - consistent with our data (Fig. 4e) - but reported to be expressed in β-cells in rats^46^. Notably, these cells also expressed progenitor-associated markers ALDH1A1 and ALDH1B1, suggesting a degree of developmental plasticity^47,48^. Additionally, the prohormone VGF, which regulates insulin granule formation and β-cell function, was enriched in this population, supporting the idea that polyhormonal cells may contribute to islet cell replenishment and adaptation^49^.

Comparison of different RBP4-positive polyhormonal cells revealed hundreds of differentially expressed proteins (Fig. 5d). Notably, NPDC1/RBP4/SST-positive cells exhibited upregulation of α-cell markers (GCG, NPDC1, GC), alongside prohormones (SCG2, VGF, CPE) and ARX, a transcription factor that directs pancreatic endocrine progenitors toward the α/PP developmental pathway. Loss of ARX has been shown to trigger the reprogramming of adult α- cells into β-cells, underscoring a potential role for these polyhormonal populations in islet plasticity and adaptive endocrine remodeling^51^. Importantly, all markers identified through multiplexed staining were detected in our proteomic analysis. Additionally, we identified the prohormone ALDH1A1, as well as PCSK2, a marker of immature endocrine cells in CPEP/RBP4/SST-positive cells^52^. In contrast, CPEP/RBP4-positive cells exhibited upregulation of CDKN1C, a cell cycle inhibitor known to be expressed in mature β-cells, where it plays a critical role in limiting proliferation.

### Proteomic insights into hybrid polyhormonal endocrine states in pancreatic islets

Next, we examined the expression patterns of key islet cell markers, progenitor-associated factors, and proliferation markers across different cell populations (Fig. 5e). As expected, α-cells predominantly expressed GCG, ALDH1B1, and GLUL, whereas β-cells showed high levels of IAPP, INS, and SCGN. Polyhormonal cells exhibited a hybrid expression pattern, co-expressing islet markers while also displaying elevated proliferation markers and SCG3 expression. Beta with SST expression cells mirrored β-cell marker expression but showed higher SST levels. Notably, SCG3 isoform 2 was more highly expressed in beta with SST expression cells than in other cell types, suggesting potential functional differences. Interestingly, both beta with SST expression and polyhormonal cells demonstrated increased expression of distinct SCG3 isoforms. Stridsberg et al.^53^ previously reported Secretogranin co-expression in adult human islet cells, but its functional significance remains unclear. More recent studies have implicated SCG3 in carbohydrate metabolism, diabetes pathogenesis, and its role as a pro-angiogenic factor in diabetic retinopathy^54^. Additionally, SCG3 expression has been linked to NKX2.2 regulation and is potentially involved in β-cell mass loss through mature β-cell dedifferentiation and diabetes progression^55^. Another notable finding was the elevated expression of INSM1 in polyhormonal and beta with SST expression cells. INSM1 is a transcription factor known to play a crucial role in islet cell differentiation, β-cell maturation, and transdifferentiation from acinar to β-cells^56,57^. However, the precise mechanisms underlying its regulation and potential interactions with other non-conventional transcription factors in islet cell maturation and disease progression remain poorly understood.

Comparison of our proteomic data with publicly available single-cell transcriptomic datasets for the same markers revealed overall concordance in the expression trends of conventional endocrine markers such as GCG, SST, IAPP, INS, and TTR (Fig. 5f)^58^. However, gene expression data failed to capture similar trends in polyhormonal cells, particularly for SCG3 (Kruskal-Wallis test, p-value 0.0012) and INSM1 (Kruskal-Wallis test, p-value 0.0233), which were detected at the protein level in our dataset.

These findings underscore the power of our mxDVP pipeline in resolving complex cellular landscapes and identifying rare cell states within highly heterogeneous human tissues such as pancreatic islets. By integrating high-resolution multiplexed imaging with Deep Visual Proteomics, mxDVP enables in-depth characterization of rare endocrine subpopulations, including polyhormonal and progenitor-like cells, while providing protein-level insights that complement and extend transcriptomic analyses. The identification of distinct prohormone expression patterns, transcriptional regulators, and proliferation-associated factors in polyhormonal cells suggests a potential role in islet cell plasticity and adaptation. Notably, proteins such as SCG3 and INSM1, which are implicated in endocrine differentiation and β-cell function, exhibited differential expression in polyhormonal and beta with SST expression hybrid cells, highlighting avenues for further investigation into islet cell fate dynamics. By offering spatially resolved proteomic data in highly heterogeneous tissues, mxDVP provides a powerful and scalable framework for studying cellular diversity across different organ systems in both health and disease. These insights contribute to a deeper understanding of islet biology and may inform future research into tissue homeostasis, regeneration, and diabetes pathophysiology.

## Discussion

Our study introduces multiplexed Deep Visual Proteomics (mxDVP) as a novel approach for investigating pancreatic islet cell heterogeneity with unprecedented spatial and molecular resolution. By integrating multiplexed imaging with ultra-sensitive mass spectrometry, mxDVP provides insights into the complex cellular architecture of pancreatic islets as well as deep proteomic profiles to understand cellular activity, revealing rare endocrine states and spatial relationships that were previously elusive.

The identification of rare polyhormonal cells in normal pancreatic tissue underscores the functional plasticity of endocrine cells, challenging traditional views of islet composition. These rare cell populations, exhibiting co-expression of markers from multiple endocrine lineages, may represent transient states or early stages of transdifferentiation, possibly as an adaptive response to metabolic demands^59^. Although the current donors show low BMI and relatively healthy insulin secretion, these findings could indicate latent plasticity that may become more pronounced under conditions of metabolic stress, such as in older individuals, those with obesity, or those in a pre-diabetic state. Indeed, a broader comparison with these populations would help validate the hypothesis that these rare cell types are involved in the islet response to stress. Such findings would align with previous studies linking cellular plasticity to β-cell dedifferentiation and highlight potential mechanisms for maintaining islet function under stress^60^. Furthermore, the discovery of distinct endocrine subtypes suggests that cellular heterogeneity within islets extends beyond classical α-, β-, and δ-cell categorizations, revealing a spectrum of cellular identities with unique metabolic and regulatory profiles^61,62^.

Spatial analysis further elucidates the organizational principles underlying islet biology. We observed that larger islets, predominantly composed of classical endocrine subtypes, exhibit reduced vascularization compared to smaller, polyhormonal cell-enriched islets. This differential vascularization may support distinct metabolic states or microenvironmental adaptations, such as hypoxic stress in larger islets. The correlation between islet size and cellular composition suggests that islet architecture may be dynamically regulated, influenced by local nutrient availability and metabolic demands^63,64^.

Proteomic profiling of endocrine subtypes revealed unique pathway enrichments, with β-cells showing enhanced metabolic pathways and α-cells enriched for hormone secretion-related pathways^65^. The presence of transcription factors associated with progenitor states in polyhormonal cells suggests these populations may contribute to islet regeneration or plasticity^66^. Moreover, the differential expression of prohormones and immune-related proteins within these cells highlights their potential roles in local immune modulation and metabolic adaptation, providing new avenues for exploring islet cell resilience in diabetes and other metabolic disorders.

Importantly, we demonstrate that even when constrained to a limited panel of gold-standard markers typically used in routine immunofluorescence or immunohistochemistry, our findings align with known endocrine cell distributions and heterogeneity trends reported in prior studies. However, the use of expanded panels incorporating additional endocrine subtype-specific markers allows more refined stratification of transient and rare endocrine states. When this refined spatial stratification is coupled with deep proteomic profiling, it enhances the resolution with which we can characterize intermediate or plastic states, offering mechanistic insights that are otherwise inaccessible through conventional marker sets alone.

Beyond pancreatic islets, the adaptable mxDVP pipeline provides a powerful framework for studying tissue heterogeneity across diverse physiological and pathological contexts. Recent applications of the standard Deep Visual Proteomics workflow have led to key translational biomedical discoveries, including the identification of novel targets in Hodgkin lymphoma, such as IRF4, proteasome subunits, and IL-signaling pathways^8^, as well as targetable JAK signaling in toxic epidermal necrolysis^7^. These studies underscore the potential of DVP to uncover disease mechanisms, inform targeted interventions, and advance precision medicine. Our approach offers several advantages over traditional methodologies, including higher spatial precision, minimal tissue requirements, and compatibility with archived tissue specimens. By utilizing a high-plex antibody panel, refined segmentation algorithms, and simplified image analysis through PIPΣX, mxDVP enables robust characterization of cell populations, ensuring that rare or transitional states are not overlooked. This methodology facilitates the direct correlation of spatial organization with molecular signatures, enhancing our understanding of how cellular microenvironments influence function and pathology.

By offering an integrated end-to-end solution for high-resolution spatial proteomics, mxDVP bridges the gap between imaging and deep molecular profiling, enabling targeted in-depth characterization of rare cell populations that would otherwise remain undetected. These findings lay the groundwork for future investigations into islet cell fate dynamics and provide a blueprint for extending spatial proteomic approaches to other complex tissues. As spatially resolved proteomics continues to evolve, its application in understanding tissue microenvironments, disease progression, and therapeutic response is poised to drive advances in regenerative medicine and precision therapeutics.

## Materials and Methods

### Human sample procurement

Formalin-fixed paraffin-embedded (FFPE) tissue samples from the pancreas tail were obtained from two female donors were provided by the Alberta Diabetes Institute (ADI) IsletCore (http://www.bcell.org/adi-isletcore.html)^67^. All donors’ families gave informed consent for the use of pancreatic tissue in research (University of Alberta Human Research Ethics Board approval Pro00013094). The research was approved by Etikprövningsmyndigheten (Dnr 2020-01690). Donors in the cohort were of similar age, of the same sex, and had no previous history of either diabetes mellitus or pancreatitis. Donor characteristics are described in Supplementary Table 1 and full phenotyping information for this tissue can be found at www.humanislets.com.

### Development of custom spatial proteomic pancreatic antibody assay

To allow visualization of key cell types and hormonal states in pancreatic tissue using multiplexed imaging, we developed and validated a panel of DNA-barcoded antibodies.

### Antibody validation by immunostaining

The targets included in the antibody panel were selected based on a combination of literature and patch-seq data^68,69^. Antibodies toward targets not supplied by Akoya Biosciences were sourced from commercial vendors as detailed in Supplementary Table 3. Clones were selected based on formulation (carrier-free), concentration (>0.5mg/mL) per conjugation kit compatibility recommendations from Akoya Biosciences (CODEX User Manual Rev C ®), and validation data supplied by vendors, ensuring that the antibodies had been validated for immunohistochemistry (IHC) in FFPE tissue.

To ensure the specificity of the antibodies to be included in the assay, each antibody was validated by indirect immunofluorescence staining, using the protocol described by Martinez Casals et. al.^70^. In brief, single sections of FFPE human, non-diabetic, adult pancreatic tissue were sectioned at 5µm thickness and mounted onto positively charged microscope slides (VWR, Epredia, Cat. No. 76406-502). The samples were deparaffinized, rehydrated, and underwent heat-induced epitope retrieval (HIER) treatment using pH 6 citrate buffer (Sigma, Cat. No. C9999- 1000mL) diluted to 1X in ddH2O in a pressure cooker (Bio SB TintoRetriever, Cat. No. BSB-7087) set to high pressure at 114–121 °C for 20 minutes. Following the HIER treatment, the samples were photobleached to reduce tissue autofluorescence. After four PBS washes, antibodies (all antibodies labeled as “Custom” in Supplementary Table 3) were diluted in antibody diluent containing 0.3 % Triton (Sigma Aldrich, Cat. No. T8787) 1X PBS and subsequently added onto the samples and incubated at 4 °C overnight. The next day, the samples were washed three times with TBS-T (1X TBS (Medicago, Cat. No. 09-7500-100) 0.1% Tween (Sigma, Cat. No. P1379) and blocked using TNB (0.5% TSA blocking reagent (Akoya Biosciences, Cat. No. FP1012) in 1X TBS (Medicago, Cat. No. 09-7500-100) for 30 min. Next, secondary antibodies and nuclear stain diluted in TNB (1:800 secondary antibody as detailed in Supplementary Table 3 and 1:400 Hoechst (ThermoFisher, Cat. No. H3570), were added to the sections and incubated for 90 minutes at room temperature. Following three washes with TBS-T, the samples were mounted with Fluoromount-G (ThermoFisher, Cat. No. 00-4958-02) prior to image acquisition.

### Custom antibody conjugation

Pre-conjugated antibodies supplied by Akoya Biosciences were purchased with their respective barcodes as detailed in Supplementary Table 3. Antibodies that had been validated were custom-conjugated to their respective barcodes as described in Supplementary Table 3 following the conjugation protocol in Akoya Biosciences’ CODEX User Manual Rev C ® using the Conjugation Kit (Akoya Biosciences, Cat. No. 7000009). In brief, 50µg of antibody was added onto a 50kDa MWCO filter (EMD Millipore, Cat. No. UFC505096) and thereafter concentrated by centrifugation. Next, the antibodies underwent a reduction step through incubation with a reduction master mix at room temperature for 30 minutes. Following the reduction step, the buffer of the antibodies was exchanged for the Conjugation Solution (Akoya Biosciences, Cat. No. 7000009) by centrifugation. Next, lyophilized barcodes were resuspended in the Conjugation Solution, added to each respective antibody, and incubated for 2 hours at room temperature. Following three washes by buffer exchange to the Purification Solution (Akoya Biosciences, Cat. No. 7000009), 100µL of Antibody Storage Solution (Akoya Biosciences, Cat. No. 7000009) was added to the filter, and the purified, conjugated antibody was eluded and stored at 4 °C.

### Antibody conjugate validation by tissue staining

To validate whether the conjugation reaction was successful, the custom-conjugated antibodies were stained individually in FFPE human, non-diabetic, adult pancreatic tissue and visualized by manual addition of the complementary fluorescently labeled oligonucleotide sequence, also referred to as a reporter, as detailed in the CODEX User Manual Rev C ® by Akoya Biosciences. Largely the same protocol for tissue staining was used as described by Martinez Casals et. al., with a few modifications to the end of the protocol to allow for the manual addition of reporters^71^. In brief, the same protocol for dewaxing, rehydration, antigen retrieval, and photobleaching was used as described above for immunostaining of tissues. Once the photobleaching was complete, the samples were washed twice in Hydration Buffer (Akoya Biosciences, Cat. No. 7000017) before being equilibrated in Staining Buffer (Akoya Biosciences, Cat. No. 7000017) for 30 minutes. Next, custom-conjugated antibodies diluted in a solution of Staining Buffer N, J, G, and S blockers (Akoya Biosciences, Cat. No. 7000017) were added to the samples and incubated at 4 °C overnight. The next day, the samples were washed twice in Staining Buffer before being fixed with 1.6% paraformaldehyde (ThermoFisher, Cat. No. 043368) diluted in Storage Buffer (Akoya Biosciences, Cat. No. 7000017) through a 10-minute incubation at room temperature. Following three washes with 1X PBS, the samples were fixed in ice-cold methanol (Sigma-Aldrich, Cat. No. 322415) for 5 minutes. After three washes with 1X PBS, the samples were fixed with Fixative Solution (Akoya Biosciences, Cat No. 7000017) diluted 1:50 in 1X PBS through a 20-minute incubation at room temperature before being washed three times with 1X PBS. Once the post-fixation was complete, the samples were washed three times with Screening Buffer (20% dimethylsulfoxide (DMSO) (Sigma-Aldrich, Cat. No. 472301) in 1X CODEX Buffer (Akoya Biosciences, Cat. No. 7000001). Next, reporters diluted 1:40 in Reporter Stock Solution (5% Assay Reagent (Akoya Biosciences, Cat. No. 7000002) and 0.25% Hoechst (ThermoFisher, Cat. No. H3570) in Screening Buffer) were added onto the samples and incubated for 5 minutes at room temperature before the samples were washed three times with Screening Buffer. Lastly, the samples were washed once with 1X CODEX Buffer (1:10 dilution of 10X CODEX Buffer (Akoya Biosciences, Cat. No. 7000001) in ddH2O) before they were mounted with Fluoromount-G before image acquisition.

### Image acquisition

Images were acquired using a Leica DMI8 widefield microscope equipped with a digital CMOS camera (ORCA-Flash 4.0 V3, Hamamatsu) and SOLA-SM-II light source at 20X magnification (20x/0.75 NA dry objective with a resolution of 0.325µm/pixel) at 16-bits through the Leica Application Suite X (LAS X, v. 3.7.5.24914). Acquired images were merged and analyzed in ImageJ (v. 1.53) to determine the signal-to-noise ratio (SNR). A cutoff of 3 was used for the SNR for both primary antibody and antibody conjugate validations. The staining of each antibody was evaluated by comparing it to reference images provided by the manufacturer and the Tissue Atlas section of the Human Protein Atlas (HPA) to validate the specificity. The developed antibody panel has previously been published as an Organ Mapping Antibody Panel (OMAP) as part of the Human BioMolecular Atlas Program (HuBMAP)^72^. A representative dataset of CODEX images in human pancreas defining the main cell types and anatomical structures of human pancreatic tissue is accessible through OMAPs^73^.

### Highly multiplexed staining and imaging on membrane slides Multiplexed immunofluorescence staining (CODEX)

For the highly multiplexed (CODEX) staining of the samples included in the cohort, single sections of FFPE human, non-diabetic, adult pancreatic tissue were sectioned at 8µm thickness and mounted onto 22mm x 22mm x 4um custom-designed metal-framed PPS membrane coverslips (Microdissect). To dewax the samples, the membrane coverslips were placed onto glass slides with 100µL ddH2O underneath and into a sealed packet of aluminum foil before being baked on a hot plate at 55 °C for 30 minutes to promote tissue adherence to the membrane and prevent excess dehydration. Once dewaxing was complete, the same protocol for CODEX tissue staining as described by Martinez Casals et. al. was followed, with the exception that no photobleaching protocol was performed^71^. After the three post-fixation steps were completed, the samples were placed in Storage Buffer (Akoya Biosciences, Cat. No. 7000017) and stored at 4 °C.

### CODEX image acquisition

Reporters were prepared according to the instructions of the CODEX User Manual Rev C ® by Akoya Biosciences and added to 96-well plates (Akoya Biosciences, Cat. No. 7000006). Prior to image acquisition, samples were equilibrated to room temperature in 1X CODEX Buffer (Akoya Biosciences, Cat. No. 7000001). The experimental template was prepared in CODEX Instrument Manager (CIM, v. 1.30.0.12), a software that automates the fluidic cycles and image acquisition, where the order of addition of reporters and exposure times were specified as detailed in Supplementary Table 4. Images were acquired with the same Leica DMI8 widefield microscope as described above at 20X magnification and 16-bit depth. To allow for visualization of the samples and CODEX image acquisition set-up, the samples were incubated with Hoechst (ThermoFisher, Cat. No. H3570) diluted 1:1000 in 1X CODEX Buffer for 3 minutes at room temperature before being washed once with 1X CODEX Buffer. A region of interest (ROI) was defined manually for each sample (ranging between 77 to 160 tiles, each tile 2048x2048 pixels and with a 10% overlap between tiles), and the focus was set automatically by LAS X for all manually placed focus points evenly distributed across the ROI. Each tile in the ROI was imaged as a 14.99 µm Z-stack, where each Z-step was 1.50 µm resulting in a total of 11 images in the Z-axis.

Images were processed and stitched in the CODEX Processor version 1.7.0.6 using background subtraction, deconvolution (vectorial, 25 iterations) and shading correction. The extended field feature was not enabled during processing.

### Tissue storage and polychromatic staining pre-laser microdissection

Post-imaging tissue preservation conditions were optimized to maintain structural integrity. Once the CODEX imaging was completed, each sample was immediately removed from the microscope and washed twice for 1 minute in ultrapure water (ThermoFisher, Cat. No. 51140) before being dehydrated through a series of 1-minute incubations in increasing concentrations of ethanol (SigmaAldrich, Cat. No. 1009832500) (30%, 50%, 70%, 80%, and 90%) and a final 2- minute incubation in 100% ethanol. Lastly, the samples were stored at 4 °C in 100% ethanol to preserve the tissues. Prior to laser capture microdissection samples were hydrated for one minute at 100%, 95% and 70% Ethanol followed by ultrapure water for 2 minutes. Finally, the sample was stained with Toluidine blue (Paradise Plus, ThermoFisher, Cat. No. KIT0312S) for 45 seconds and washed for 10 seconds with 75% ethanol before being loaded on the LMD7 system. A pseudocoverslipping step, involving the addition of 10 µL of 70% ethanol to the tissue section before imaging, enabled high-quality reference image acquisition for precise contour import. The tissue sections were dried for 2 minutes before dissecting contours. Microdissected samples were collected directly into a protein low-bind 384-well plate and stored dry at 4 °C after cutting.

### Integrated image analysis pipeline for segmentation and cell annotation

We centralized the image analysis pipeline by implementing an in-house developed software, PIPΣX, which allows performing all image processing steps i.e. pre-processing, segmentation, post-processing, and analysis in one place^11^. TIFF images were uploaded to the PIPΣX web platform (version 1.0) for automated processing. Segmentation, filtration, and clustering analyses were performed using the following parameters: Nuclear segmentation was conducted with Hoechst staining, applying a nucleus diameter of 20 pixels (equal to 6.5 µm) and a definition threshold of 0.5 via the integrated StarDist algorithm. Membrane refinement was performed following nuclear segmentation, leveraging CDH1 intensity. The refinement parameters were set to a membrane diameter of 25 pixels (equal to 8.125 µm) and a compactness value of 0.9. The membrane refinement confidence score, calculated as the proportion of cell contours ultimately defined by CDH1 membrane staining during watershed expansion, reached 96%. PIPΣX generated quantification output as a CSV table, reporting marker intensities as the mean pixel intensity per region of interest (each cell contour). These steps were executed using a locally installed beta version of PIPΣX. To remove artifacts, cells were filtered based on size, excluding the bottom 1% and top 5% of the size distribution.

#### Semi-automated clustering analysis

First, K-means clustering was applied to non-normalized data, with K = 25, for each tissue section separately. A subsequent cluster refinement algorithm was implemented, incorporating ranking scores for each marker. The algorithm evaluates each cluster by comparing its ranked markers to predefined rules (annotations), assigning a confidence score based on alignment. In the final refinement step, the cluster annotation with the highest confidence is selected. Only markers with expression levels above zero were considered for refinement.

#### Clustering of endocrine cell populations

To annotate endocrine cell populations, output tables from each tissue section were merged and normalized using log1p transformation and quantile normalization within PIPΣX (Supplementary Table 5). Endocrine cells were subsetted and further stratified by tissue section before applying K-means clustering to the normalized data. Various K-values were tested, with K=7 yielding the most robust results. Clusters were annotated based on Spearman rank correlation scores, calculated using only endocrine cell marker intensities. This approach identified 12 distinct endocrine cell subpopulations with diverse marker expression profiles.

To validate segmentation accuracy and cell annotation, the annotated clusters were exported to the PIPΣX-integrated TissUUmaps software for visualization, facilitating dynamic exploration of annotations, cellular neighborhoods, spatially reduced data, and segmentation masks^74,75^. Segmentation masks and cluster assignments were reviewed against the original images to ensure fidelity.

### Spatial analysis

The integrated dataset of annotated cell populations was used to analyze spatial associations. Mean intensities for each cell population were calculated, followed by Euclidean distance analysis to assess spatial relationships across all donors and annotated cell types.

To further investigate the spatial organization, we used TissUUmaps to visualize the spatial neighborhoods and quantify islet sizes with the Eclipse Drawing plugin. Islet diameters were extracted from the detailed report generated by TissUUmaps. To examine the immediate microenvironment, we expanded the islet perimeter by a factor of 1.5 and performed repeated measurements within the adjusted boundary. To assess associations between islet size and cellular composition, we performed a Chi-square test using the scipy.stats library in Jupyter Notebook (Python 3.9.21).

### Segmentation mask creation and processing for BIAS integration

Cell segmentation masks were generated by PIPΣX as TIFF files. For integration into BIAS software, each mask was tiled into 2048-pixel sections without overlap. A modified PIPΣX tiling feature ensured that the final tile in each row was extended to accommodate non-standard dimensions. The number of resulting tiles varied based on the original mask size.

### Contour processing in BIAS

Image files of Hoechst staining and tiled segmentation masks were uploaded into the software BIAS (Version 1.1.0, Single-Cell Technologies Ltd.). A data frame based on the imported segmentation mask was generated in BIAS containing cell area as well as x and y coordinates and exported to be aligned with the PIPΣX data frame matching cell IDss and incorporating an additional column containing the cluster number for each ID. The modified data frame was reimported into BIAS and clusters were selected individually to process contours. We dilated cell masks by half of the size of the laser impact line (3 µm) to improve target precision and merged (melded) some of the contours based on their spatial landscape to avoid sample loss (Extended Data Fig. 2c). The contours were modified by melding touching contours, closing sharp edges at 5µm and expanding contour borders by 3µm. The melding feature allowed to minimize the subsequent sample isolation time on the LMD.

Alignment points were placed in BIAS on peripheral nuclei and exported as an xml file containing cluster contour coordinates.

### Laser microdissection

For contour isolation, we used the Leica LMD7 system with software version 8.3.745. All contours were excised with a 63x objective and collected in low binding 384 well plates (Eppendorf, Cat. No. 0030129547) at a temperature range from 19°C to 23°C.

Power and Aperture was adjusted dynamically in the given range based on tissue topography and contour size, while larger contours were cut with increased Power and Aperture. Pulse frequency was decreased to 300 Hz when humidity reached below 30%. See Supplementary Table 6 for detailed laser settings. After each collection, contours were centrifuged at 2000 x g for 2minutes.

### Sample Preparation for MS

Peptides were semi-automatically prepared on a Bravo pipetting robot (Agilent), similar to what was described previously^76–78^. Briefly, extracted contours were centrifuged at 2000 x g for 2minutes, washed down with 7µL 100% acetonitrile on each side of the well (in total 28µL) and dried in a Vacuum concentrator (SpeedVac) at 45°C for 20 minutes. Wells were visually quality controlled. In the following incubation steps, plates were tightly sealed with two stacked aluminum lids to avoid evaporation (Thermo Fisher Scientific, Cat. No. AB0626). The contours were resuspended in 6µL lysis buffer (60 mM triethylammonium bicarbonate buffer (pH 8.5, Sigma) with 0.013% DDM (Sigma)), and cooked for 60 minutes at 95°C in a 384-well thermal cycler (Eppendorf Mastercycler) at a lid temperature of 110°C. After the addition of 1 µL of 80% ACN (final concentration 10%), samples were incubated for 60 minutes at 75°C and cooled briefly. Proteins were digested overnight at 37°C using 4 ng LysC and 6 ng trypsin (1μL in lysis buffer) at a lid temperature of 50°C. The next day, digestion was stopped by adding trifluoroacetic acid (TFA, final concentration 1% v/v). Samples were loaded onto prepared Evotip Pure (Evosep) using the Bravo robot (Agilent).

### Peptide loading onto C-18 tips

C18 tips (Evotip Pure, Evosep) were activated in 1-propanol, washed two times with 50µL buffer B (99.9% ACN, 0.1% FA), activated again in 1-propanol and washed twice with 50µL buffer A (99.9% H2O, 0.1% FA). In between Evotips were spun at 700 g for 1 minute. Evotips were prepared with 70µL buffer A and a short spin at 700 x g for sample loading. Samples were loaded into the remaining buffer A on the tip. Evotips were spun at 700 x g for 1 minute to load the sample, washed with 50µL buffer A and stored at 4°C with buffer A on top until liquid chromatography–mass spectrometry (LC–MS) analysis.

### LC-MS/MS analysis

Samples were loaded onto Evotips Pure and measured with the Evosep One LC system (Evosep) coupled to an Orbitrap Astral mass spectrometer (Thermo Fisher Scientific). The Whisper40 SPD (samples per day) method was used on a commercial analytical column Aurora Elite TS of 15cm length and 75 μm inner diameter (ID), packed with 1.7 μm C18 beads (IonOpticks). The column temperature was maintained at 50°C using a column heater (IonOpticks). The mobile phases comprised 0.1% FA in LC–MS-grade water as buffer A and 99.9% ACN/0.1% FA as buffer B. The Orbitrap Astral mass spectrometer was equipped with a FAIMS Pro interface and an EASY-Spray source (both Thermo Fisher Scientific) at electrospray voltage of 1900V. A FAIMS compensation

voltage of -40V and a total carrier gas flow of 3.5 L/min were used. The Orbitrap Astral was operated in DIA (data independent acquisition) mode with variable window widths at a positive spray voltage of 1900 V. The Orbitrap analyzer was utilized for full MS1 analyses with a resolution setting of 240,000 within a full scan range of 380–980 m/z. The automatic gain control (AGC) for the full MS1 was adjusted to 500% with a maximum injection time of 100 ms. The Astral analyzer was used for MS/MS acquisition with isolation windows of 6 Th (100 windows), a maximum ion injection time (IIT) of 10 ms, an AGC target of 800%, and the MS/MS scanning range of 150−2000 m/z. Selected ions were fragmented by higher-energy collisional dissociation (HCD) at a normalized collision energy (NCE) of 25%.

The quality control samples were measured with the Evosep One LC system (Evosep) coupled to a timsTOF SCP mass spectrometer (Bruker). The Whisper20 SPD (samples per day) method was used on an Aurora Elite CSI third generation column with 15 and 75 μm ID (AUR3-15075C18-CSI, IonOpticks). All other chromatography settings were the same as described above. The timsTOF SCP was operated in dia-PASEF mode with variable window widths. Optimal dia-PASEF methods cover the precursor cloud highly efficient in the m/z – ion mobility (IM) plane while providing deep proteome coverage. The optimal dia-PASEF method generated with py_diAID^79^ consisted of one MS1 scan followed by twelve dia-PASEF scans with two IM ramps per dia-PASEF scan, covering a m/z range from 350 to 1,200 and IM of 0.7 to 1.3 Vs cm^−2^ ^77^. The mass spectrometer was operated in high sensitivity mode, the accumulation and ramp time was specified as 100 ms, capillary voltage was set to 900 V and the collision energy was a linear ramp from 20 eV at 1/K0 = 0.6 Vs cm^−2^ to 59 eV at 1/K0 = 1.6 Vs cm^−2^.

### Raw data analysis with DIA-NN

Orbitrap Astral raw files were first converted to the mzML file format using the MSConvert software (https://proteowizard.sourceforge.io/) from Proteowizard, keeping the default parameters and selecting ‘Peak Picking’ as filter. Raw files (mzML or Bruker .d) were searched using the DIA-NN software^80,81^ (version 1.8.1) using the FASTA (2023, UP000005640_9606, with 20,594 gene entries) from the UniProt database and a library-free approach. A deep-learning module, match-between-runs (MBR) and heuristic protein inference (‘--relaxed-prot-inf’) was enabled. N-terminal methionine excision was set as fixed modification and oxidation as variable with maximum one allowed. ‘IDs, RT & IM profiling’ was used for library generation, ‘robust LC (high accuracy)’ for quantification and ‘RT-dependent’ for cross-run normalization. One miss cleavage was allowed and precursor charge range was set from 2 to 4. The false discovery rate (FDR) was controlled at less than 1% for peptide spectrum matches and protein group identifications. All other parameters were set as default. The ‘pg_matrix.tsv’ output file was used for further data analysis (Supplementary Table 7).

### Bioinformatic data analysis

All statistical analyses were performed using Perseus (v2.0.10), Python (v3.9.4), and R (v4.2.2). Initial data processing included filtering out proteins from common contaminants, followed by log2 transformation.

For differential expression analysis, Welch’s t-test with a permutation-based false discovery rate (FDR) of 5% was conducted in Perseus. Enrichment analyses were performed using Fisher’s exact test with an FDR cutoff of 0.02, using the full dataset of 6,391 proteins as background. Supervised subtype-based analysis was performed using one-way ANOVA followed by post hoc tests, with a permutation-based FDR cutoff of 0.05 in Perseus. No imputation was applied for supervised comparisons.

Data visualization was carried out in Python using the seaborn package. Protein interaction networks were constructed using STRING output and visualized in Cytoscape (v3.8 and v3.10.3). One-dimensional (1D) annotation enrichment analysis was performed on score values using Perseus, applying a Benjamini–Hochberg FDR threshold of 0.05.

For principal component analysis (PCA), samples with fewer than 4000 identified proteins were excluded. Missing values were imputed using a normal distribution, downshifted by 1.9 standard deviations with a width of 0.5 relative to the original distribution.

Comparative analyses with annotated single-cell RNA sequencing (scRNA-seq) data were performed using datasets from Soo Yoon et al.^82^ and Tabula Sapiens^83^.

## Code and Data Availability

The high-plex image data is available in EBI Image Archive BioImages^84^ (https://www.ebi.ac.uk/bioimage-archive/). All mass spectrometry raw files and proteomics data analyzed in this study have been deposited to the ProteomeXchange Consortium via the PRIDE partner repository^85^ (http://www.ebi.ac.uk/pride) and will be made publicly available upon publication. All code for image preprocessing, segmentation, postprocessing and analysis associated with current submission is provided as a part of PIPΣX. It is available at https://github.com/CellProfiling/pipex. The PIPΣX-BIAS integration script is available at https://github.com/CellProfiling/ell_code_template/blob/master/project_utils/MXDVP/bias_pipex_match.py

## Acknowledgements

We gratefully acknowledge funding from HIRN (NIH 1U01DK120447; P.E.M., E.L), The Chan Zuckerberg Initiative (E.L.; P.H.; M.M.) EU Horizon 2020 programme (ESPACE 874710; E.L.), SciLifeLab TDP funding (E.L) and European Union’s Horizon 2020 research and innovation program under grant agreement No. 874839 (ISLET; M.T.). We also acknowledge support from the BioImage Informatics Facility (NBIS Sweden), with funding from SciLifeLab and the National Microscopy Infrastructure NMI (VR-RFI 2019-00217).

Human pancreas for research was provided by the Alberta Diabetes Institute IsletCore at the University of Alberta in Edmonton (http://www.bcell.org/adi-isletcore.html) with the assistance of the Human Organ Procurement and Exchange (HOPE) program, Trillium Gift of Life Network (TGLN), and other Canadian organ procurement organizations. Pancreas processing was approved by the Human Research Ethics Board at the University of Alberta (Pro00013094). All donors’ families gave informed consent for the use of pancreatic tissue in research.

## Author contributions

MM, ND, and EL conceived the project and designed the study. JMF and PEM collected pancreas tissue samples. MM, ND, FB, MT and AMC conducted experiments. MM, FB, MT and ND performed data analysis. MM, ND and MT curated data. FBN, FK and PH developed software. MM, ND, FB, MT and EL wrote the paper. All authors edited and commented on the manuscript.

---

**Extended Data Fig. 1:**
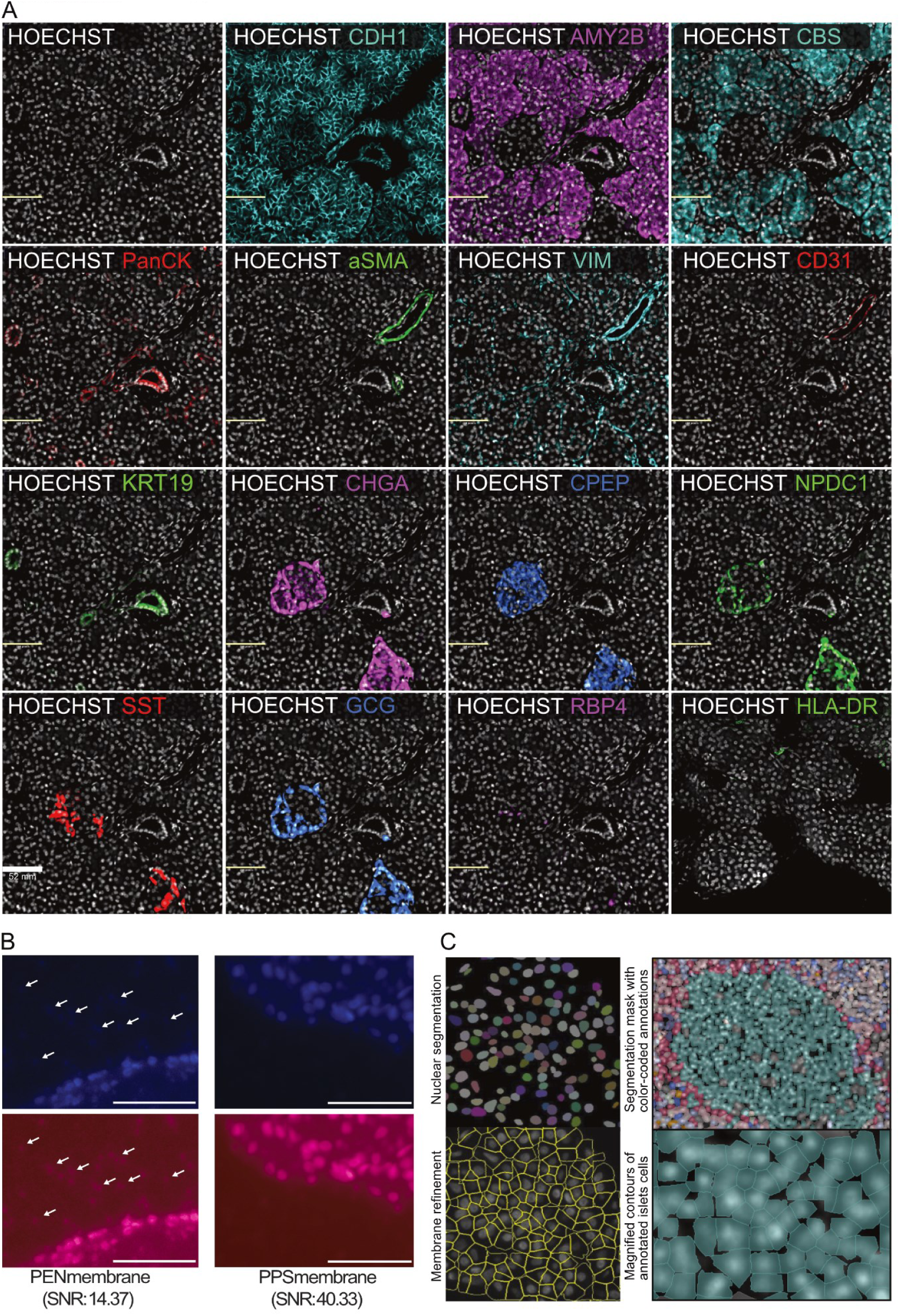
Optimization of imaging and segmentation for high-dimensional analysis of pancreatic tissue. a) Representative high-plex images from the custom-developed antibody panel showcasing the immunohistological staining of the markers of different islet cells and surrounding cellular components of the pancreatic tissue microenvironment. b) Representative images from the comparison of the image quality between untreated PEN and PPS membranes. The top panel shows the original nuclear stain (Hoechst), and the bottom panel the nuclear stain is visualized by a red Look up Table for better contrast. The white arrows indicate granules from the nuclear stain observed in PEN membranes. The scale bar is 50 µm. c) Example images from the in-house developed image analysis pipeline (PIPΣX), showcasing the nuclear segmentation by Stardist, membrane refinement feature for improved segmentation accuracy, and the full cell segmentation masks for a representative islet.

**Extended Data Fig. 2:**
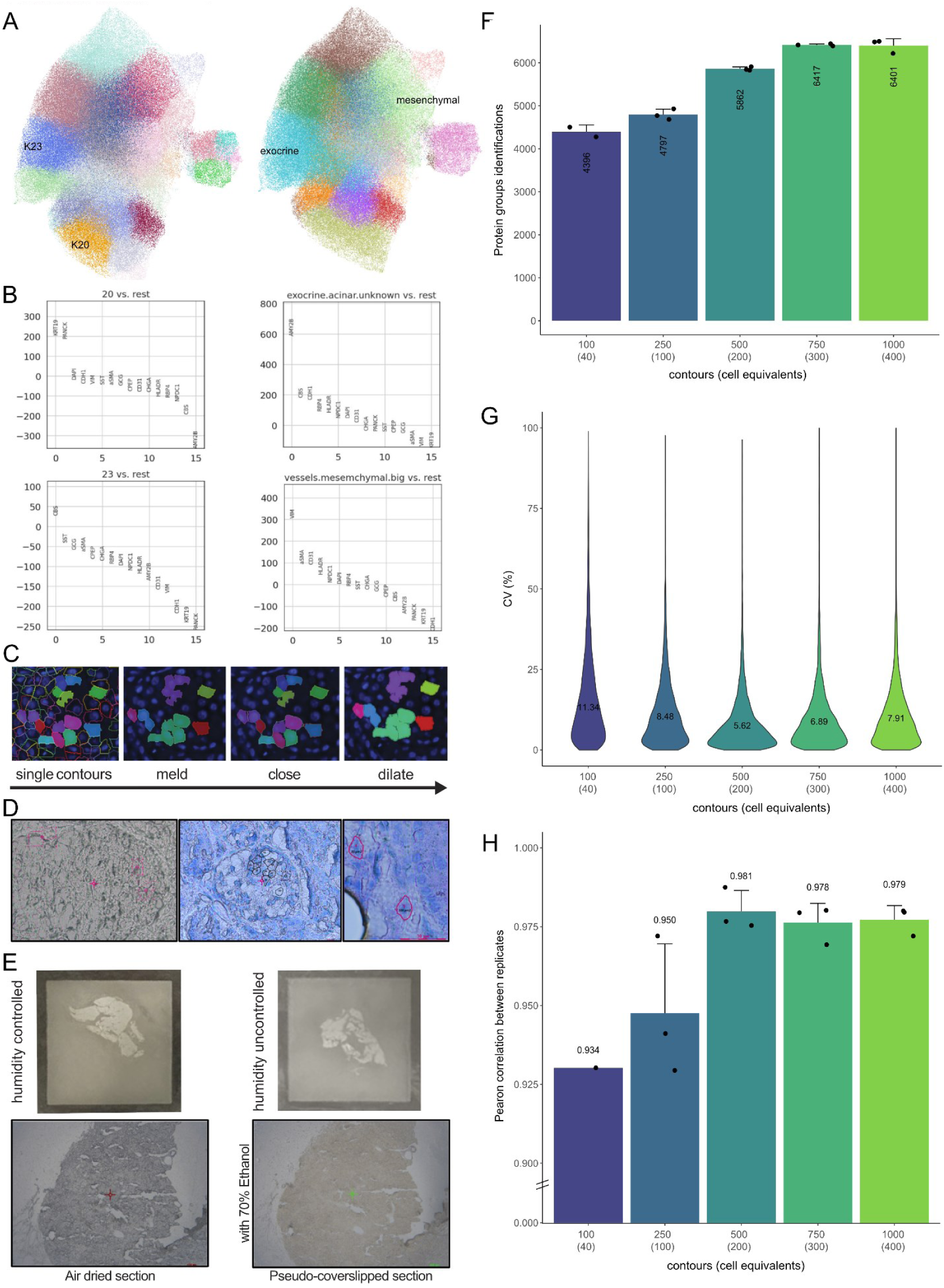
PIPΣX-enabled clustering refinement, contour optimization, and workflow validation for spatial proteomics. a) Example of a UMAP representation of all cells in one sample colored by K-means clustering with clusters 20 and 23 labeled (left). The UMAP is colored by cluster after cluster refinement in PIPΣX with the annotated cell type labeled (right). b) Corresponding plots showcasing ranked protein expression for clusters 20 and 23 before cluster refinement (left) and after cluster refinement (right). c) Representative images showcasing the developed post- adjustment tool where the exported cell segmentation masks are (1) melded based on assigned identified cell type and spatial organization, (2) gaps between the melded cell contours are closed, and (3) dilated to account for the laser impact. d) Example brightfield images with imported contours before polychromatic Toluidine stain (left) and after (right). e) Example images of coverslip PPS membranes with pancreas tissue section with controlled humidity during storage before and during tissue processing versus non-controlled humidity (left). Dry section before cutting versus pseudo coverslipped tissue with 10ul of 70% Ethanol added (right). f) Protein group identification of contour dilution series. Bar plot represents the mean of replicates, while the error bars are one standard error of the mean. Workflow replicates, n=3. g) Coefficient of variations (CVs) between replicates of contour dilution series. Workflow replicates, n=2. h) Pearson correlation between replicates of the contour dilution series. Workflow replicates, n=3.

**Extended Data Fig. 3:**
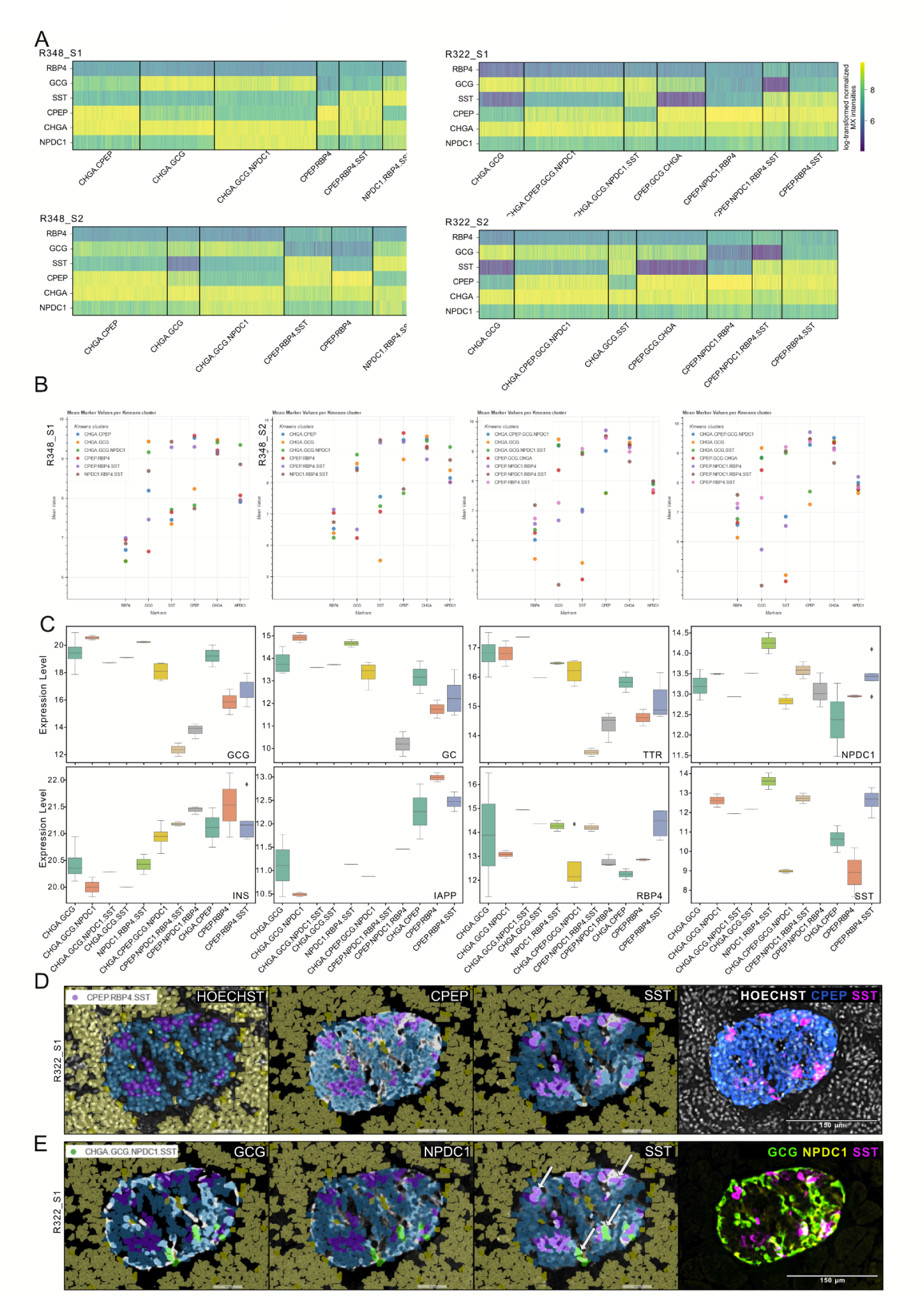
Proteomic and spatial analysis reveal endocrine cell state heterogeneity. a) Heatmaps for each donor and section displaying expression profiles of annotated islet subpopulations as identified by K-means clustering of multiplex image data (log1p-transformed normalized intensity). Multiplexed dataset was used for the comparison. b) Scatter plots for each donor and section showing the mean marker value (log1p- transformed normalized intensity) for each annotated cluster identified by K-means clustering across all islet cell markers in the antibody panel. Multiplexed dataset was used for the comparison. c) Mass spectrometry-based proteomic analysis of pancreatic endocrine cell subpopulations. The bar plot displays log2-transformed median expression levels of key endocrine markers (INS, GCG, RBP4, TTR, GC, CHGA, IAPP, SST, INSM1, NPDC1) across distinct cell subgroups. Error bars indicate standard deviations. d) Illustrative example of a CPEP-, RBP4-, and SST- coexpressing cell population. Cell segmentation masks are overlaid, with the coexpressing population shown in purple, endocrine cells in blue, and other cell types in yellow. A multiplex composite image visualizes islet marker expression within the same islet. e) Illustrative example of a GCG-, NPDC1-, and SST-coexpressing cell population within the same islet. Cell segmentation masks are overlaid, with the coexpressing polyhormonal population shown in green, endocrine cells in blue, and other cell types in yellow. An example of segmentation precision to avoid signal contamination or bleedthrough is indicated by the arrows. A multiplex composite image visualizes islet marker expression within the same islet.

**Extended Data Fig. 4:**
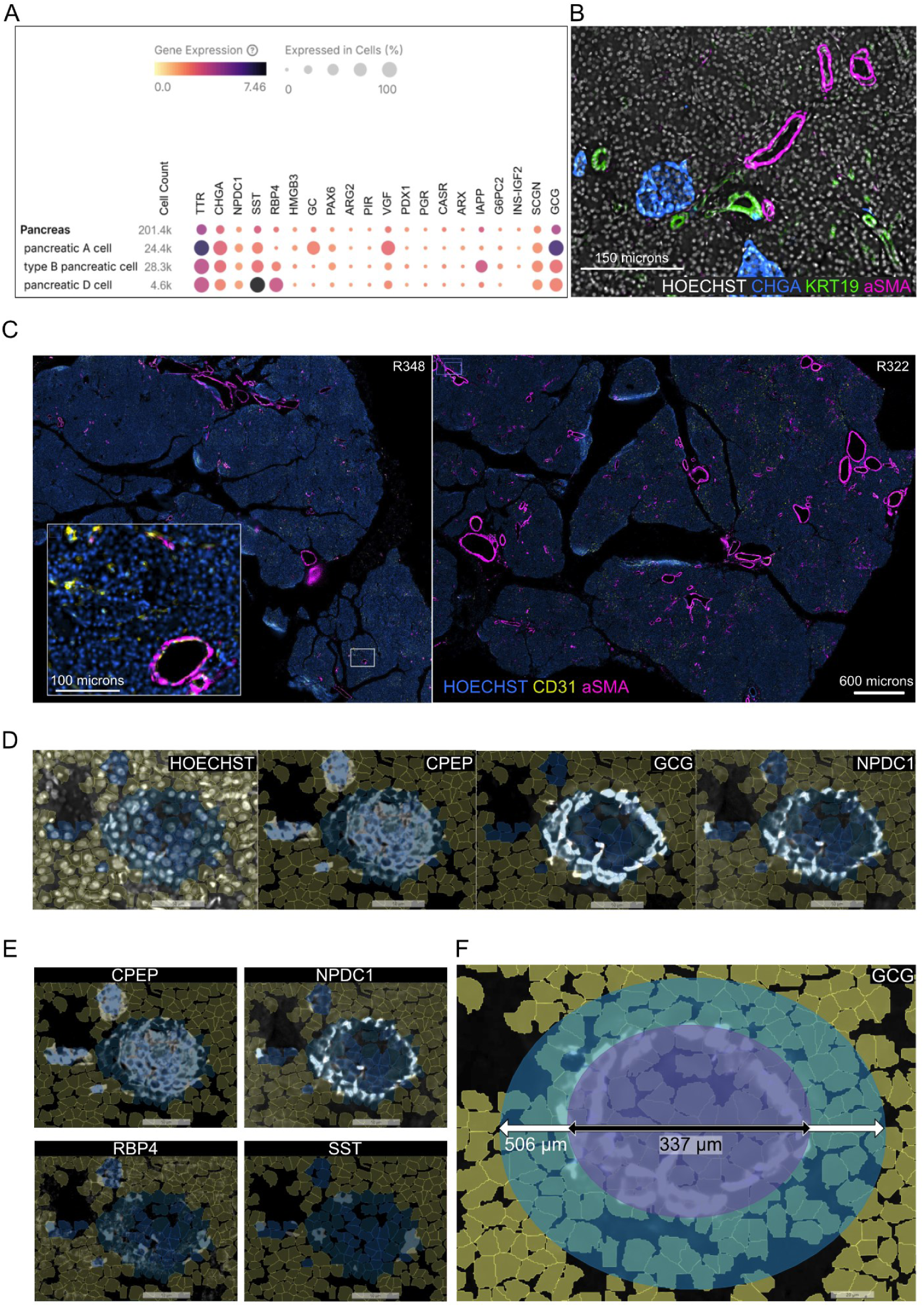
Spatial organization and compartmentalization of pancreatic endocrine cell types. a) Dot-plot highlighting differentially expressed genes for pancreatic α-cells, β-cells, and δ-cells from a publicly available single-cell annotated transcriptomic dataset (Tabula Sapiens). b) Representative high-plex image showing the observed spatially distinct organization of endocrine, ductal, and vascular compartments. c) Example multiplex images visualizing vascular markers, showcasing observed expression differences between vessels of different calibers. d) Illustrative example images of observed compartmentalization of β-cells (CPEP) and α-cells (GCG). Cell segmentation masks overlaid the images showing endocrine cells (blue) and other cell types (yellow). e) Example images of observed local cell clusters and dispersed cells of different endocrine cell populations. Cell segmentation masks overlaid the images showing endocrine cells (blue) and other cell types (yellow). NPDC1^+^RBP4^+^SST^+^ cells are showcased. f) Representative image of the investigated peri-islet microenvironment. The central ellipse superimposed on the image (purple) highlights the investigated islet, and the surrounding ellipse (blue) demarcates the peri-islet microenvironment.

**Extended Data Fig. 5:**
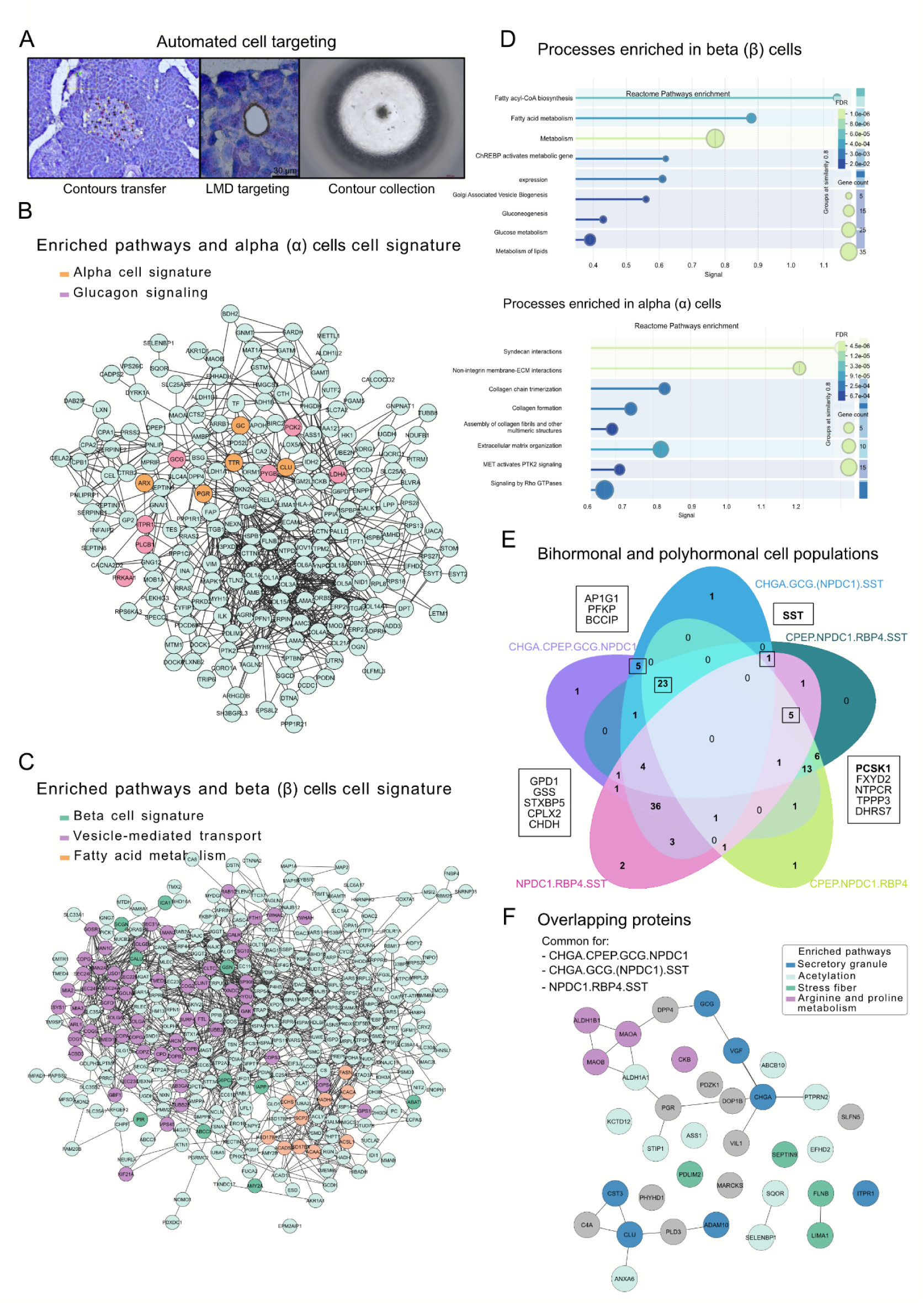
Pathway enrichment and functional profiling of rare endocrine cell populations. a) Illustrative images of key steps in the developed pipeline for automated cell targeting by LMD where (1) contours for each identified subpopulation are imported into the LMD software, (2) contours are targeted by LMD, and (3) excised contours are collected into a 384-well plate and visually quality controlled. b) Pathway enrichment analysis of α-cell proteomic expression compared to all other endocrine cells was performed using the STRING database, and pathways were colored based on enriched functions (Welch t-test, followed by Fisher exact test Benjamini-Hochberg FDR q-value < 0.02). c) Pathway enrichment analysis of β-cell proteomic expression compared to all other endocrine cells was performed using the STRING database, and pathways were colored based on enriched functions (Welch t- test, followed by Fisher exact test Benjamini-Hochberg, FDR q-value < 0.02). d) Reactome enrichment analysis of cellular processes enriched in β-cells (top) and α-cells (bottom). Statistical significance determined by FDR correction > 0.05. e) Venn diagram illustrating overlapping proteins detected by MS in isolated bihormonal and polyhormonal cell populations, with statistical significance assessed using ANOVA (p-value > 0.05), followed by post-hoc Tukey’s HSD test (FDR > 0.05) f) Enrichment analysis of the 23 proteins commonly detected in NPDC1-positive polyhormonal cells was performed and visualized based on pathway association using STRING database.

